# The actin assembly requirements of the formin Fus1 to build the fusion focus

**DOI:** 10.1101/2022.03.24.485616

**Authors:** Ingrid Billault-Chaumartin, Laetitia Michon, Caitlin A Anderson, Sarah E Yde, Cristian Suarez, Justyna Iwaszkiewicz, Vincent Zoete, David R Kovar, Sophie G Martin

## Abstract

Formins form the largest family of actin filament nucleators and elongators, involved in the assembly of diverse actin structures. Actin filament nucleation and elongation activities reside in the formin homology 1 (FH1) and FH2 domains, common to all formins. However, the rate of these reactions varies between formins by at least 20-fold. Typically, each cell expresses several distinct formins, each contributing to the assembly of one or several actin structures, raising the question of what confers each formin its specificity. Here, using the formin Fus1 in the fission yeast *Schizosaccharomyces pombe*, we systematically probed the importance of formin nucleation and elongation rates for function *in vivo*. Fus1 assembles the actin fusion focus, an aster-like structure of actin filaments at the contact site between gametes, necessary for the process of cell fusion to form the zygote during sexual reproduction. By constructing chimeric formins with combinations of FH1 and FH2 domains previously characterized *in vitro*, we establish that changes in formin nucleation and elongation rates have direct consequences on the architecture of the fusion focus, and that Fus1 native high nucleation and low elongation rates are optimal for fusion focus assembly. We further describe a point mutant in the Fus1 FH2 domain that preserves native nucleation and elongation rates *in vitro* but alters function *in vivo*, indicating an additional property of the FH2 domain. Thus, rates of actin assembly are tailored for assembly of specific actin structures.

## Introduction

Formins are a large family of conserved proteins that act as both nucleators and elongators of linear actin filaments (Breitsprecher & Goode, 2013; Courtemanche, 2018). They are involved in the assembly of many actin structures that underlie cellular processes such as polarization, motility, or division (Bohnert, Willet, et al., 2013; Goode & Eck, 2007; Pollard & O’Shaughnessy, 2019; Skau & Waterman, 2015). Most cells and organisms express different formin genes, which contribute to the assembly of distinct actin structures. For example, mammals have 15 formins (Schönichen & Geyer, 2010) and *Arabidopsis thaliana* has at least 21 (Blanchoin & Staiger, 2010), while the yeast *Saccharomyces cerevisiae* possesses only 2 (Breitsprecher & Goode, 2013) and *Schizosaccharomyces pombe* (*S. pombe*) only 3, which organize distinct actin networks (Kovar et al., 2011). In this organism, For3 supports the formation of polarizing actin cables (Feierbach & Chang, 2001), Cdc12 assembles the contractile cytokinetic ring (Chang et al., 1997), and Fus1 supports the formation of the fusion focus (Dudin et al., 2015; Petersen et al., 1995), an aster of actin filaments necessary for the fusion of gametes to form the diploid zygote. Thus, one important question is what defines the functional specificity of each formin to assemble a specific actin structure.

The defining feature of formins is their highly conserved Formin Homology 2 (FH2) domain, which forms a dimer that nucleates new actin filaments and processively binds the elongating actin filament barbed end, protecting it from growth arrest by capping proteins (Courtemanche, 2018). The FH2 domain is flanked by a FH1 domain, which extrudes from the FH2 dimer as variable numbers of flexible proline-rich tracks that bind profilin-actin to feed it to the elongating filament. The number and quality of the tracks, but also their spacing in relationship to the FH2 dimer dictate filament elongation speed (Courtemanche & Pollard, 2012; Paul & Pollard, 2008; Scott et al., 2011). For example, the fission yeast formin Fus1 possesses a single proline-rich track, compared to the 2 and 4 tracks in Cdc12 and For3 FH1 domains respectively, making Fus1 a slower elongator than these two other formins *in vitro* (Scott et al., 2011). However, because Cdc12 and For3 have very similar elongation speeds, properties other than the number of proline-rich tracks also regulate elongation speed. Furthermore, exchanging the FH1 domains between Fus1 and Cdc12 is not sufficient to convert their elongation speeds, indicating that the FH2 domain also contributes in setting elongation speed. The contribution of the FH2 domains depends on the time the domain spends in the open state (Aydin et al., 2018; Zweifel & Courtemanche, 2020). Elongation speed can also be influenced by tensile and compressive forces (Zimmermann & Kovar, 2019). In addition to F-actin nucleation and elongation, some formins exhibit additional non-canonical actin regulatory properties, contained in the FH2 or additional domains, such as actin filament bundling or severing (Courtemanche, 2018). For instance, both Fus1 and Cdc12 were shown to bundle filaments: through the FH1-FH2 domain for Fus1 (Scott et al., 2011) and through the long C-terminal tail for Cdc12 (Bohnert, Grzegorzewska, et al., 2013).

The functional specificity of diverse formins in the same cytosol is controlled in part by regulation of their localization and activation (Breitsprecher & Goode, 2013; Chesarone et al., 2010; Yonetani et al., 2008). Indeed, the FH1-FH2 domains are flanked by less conserved N- and C-terminal regulatory regions, which control formin localization and activation as well as nucleation and processivity, often through interaction with additional proteins (Breitsprecher & Goode, 2013). For instance, in Diaphanous-related formins including *S. pombe* For3, N- to C-terminal binding mediates auto-inhibition, relieved by small GTPase binding (Kühn & Geyer, 2014; Martin et al., 2007). However, many formins, such as Cdc12 or Fus1, are thought not to be autoinhibited by a canonical N- to C-terminus interaction (Scott et al., 2011; Yonetani et al., 2008). There is less information on how formins’ specific actin assembly and regulatory properties, which vary significantly between formins, participate in functional specificity *in vivo*. Recent findings by (Homa et al., 2021), have shed some light on the importance of Cdc12’s specific actin assembly properties in supporting cytokinesis, but these findings have not been generalized. The FH1-FH2 domains of all three *S. pombe* formins have been characterized *in vitro*, and their specific actin assembly rates are known (Scott et al., 2011), which makes *S. pombe* a good model organism to understand the actin assembly specificities of each formin necessary to organize its own actin network. Of interest for this work, Fus1 was shown to be a potent nucleator (1 filament per 2 formin dimers), but a modest elongator (5 subunits.s^-1^.µM^-1^).

Cell-cell fusion is a fundamental process that relies on actin assembly (Martin, 2016). In *S. pombe*, cell fusion takes place between haploid cells to form the diploid zygote during sexual reproduction, which is initiated by nitrogen starvation. Yeast cells are encased in a cell wall that protects them from external insult and internal turgor pressure. During cell fusion, the cell wall is locally digested precisely at the site of partner cell contact, to allow plasma membrane merging without risk of cell lysis. The process is coordinated by the fusion focus, an actin structure assembled by Fus1 (Dudin et al., 2015). From light and electron microscopy data, the fusion focus may be best described as an aster-like arrangement of actin filaments that concentrates glucanase-loaded secretory vesicles transported by Myosin V Myo52 promoting local cell wall digestion (Dudin et al., 2015; Muriel et al., 2021). The fusion focus shares characteristics with other formin-assembled structures: it requires profilin (Petersen et al., 1998a) and tropomyosin (Dudin et al., 2017; Kurahashi et al., 2002), and Fus1 competes with capping protein (Billault-Chaumartin & Martin, 2019). Importantly, deletion of *fus1* completely blocks cell fusion (Petersen et al., 1995). As this formin is only expressed during sexual differentiation and plays no role during mitotic growth (Petersen et al., 1995), this allows to change Fus1 properties at will and study the resulting effect without impact on viability.

In this work, we address the importance of Fus1 actin assembly properties. By systematically changing the nucleation and elongation rates through chimeras of the FH1 and FH2 domains previously characterized *in vitro* (Scott et al., 2011), we demonstrate that changes in actin assembly measured *in vitro* have direct, visible consequences on the assembly of the fusion focus, and that Fus1 actin assembly properties are tailored to its function. We further establish that the Fus1 FH2 domain contains an additional, non-canonical function that contributes to fusion focus assembly.

## Results

### Fus1 has essential properties for fusion not contained in the other *pombe* formins

To investigate whether Fus1 formin contains any unique property, we tested the ability of either of the two other *pombe* formins, Cdc12 and For3, to rescue the fusion defect of *fus1Δ* cells (Figures 1A-C). While a construct expressing *fus1* at the *ura4* locus under the *fus1* promotor in a *fus1Δ* strain was able to sustain cell fusion in a manner indistinguishable from the WT strain, constructs expressing either *for3* or *cdc12* were completely fusion deficient (Figure 1B-C). The inability of For3 and Cdc12 to complement *fus1Δ* may be due to their different N-terminal regulatory regions, which mediate localization. However, lack of localization is not the sole reason for lack of function, as For3, like Fus1, localizes to the region of cell-cell contact (Figure 1B).

**Figure 1.**
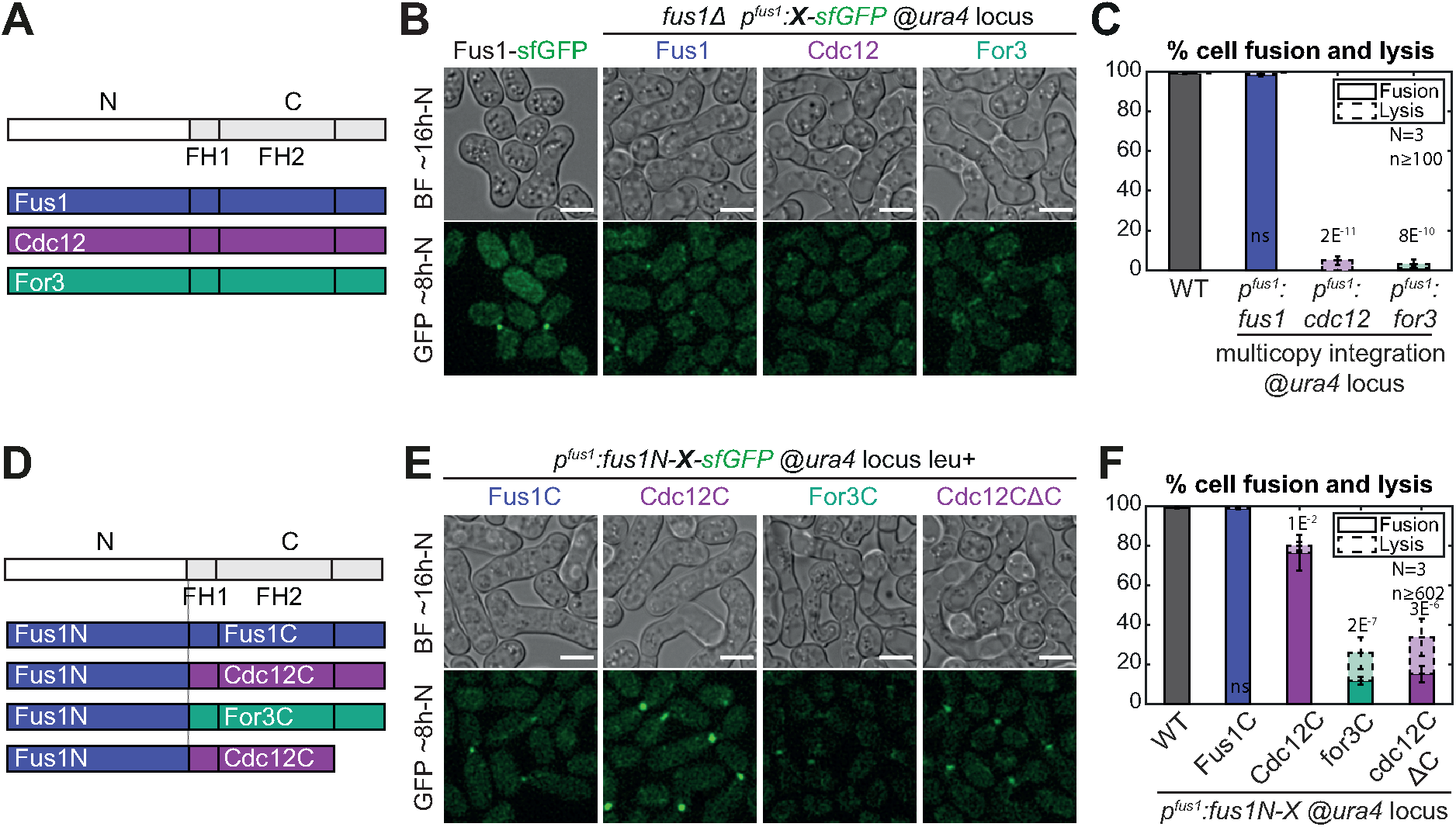
Fus1 has essential properties for cell fusion not contained in the other *S. pombe* formins. **A.** Scheme of the constructs used in panels (B-C). All are tagged C-terminally with sfGFP and expressed from the *fus1* promoter at the *ura4* locus. **B.** DIC images ∼16h post starvation and GFP fluorescence images ∼8h post starvation of homothallic WT strains expressing Fus1-sfGFP at the endogenous locus or homothallic *fus1Δ* strains expressing the formins listed in (A). **C.** Percentage of cell pair fusion and lysis 24h after nitrogen removal in strains as in (B). **D.** Scheme of the constructs used in panels (E-F). Formins are cut between the N-terminal regulatory region and FH1 domain with a BamHI restriction site, tagged C-terminally with sfGFP and expressed from the *fus1* promoter. **E.** DIC images ∼16h post starvation and GFP fluorescence images ∼8h post starvation of homothallic *fus1Δ* strains expressing the formin chimeras described in (D) at the *ura4* locus. **F.** Percentage of cell pair fusion and lysis 24h after nitrogen removal in strains as in (E) compared to the WT. Error bars are standard deviations. All p-values (student’s t-test) are relative to WT. Bars are 5µm.

Nevertheless, to alleviate any problem that would stem from improper localization or lack of other regulatory elements, we constructed a set of chimeric formins that keep Fus1N constant while varying the C-terminal part, similarly expressed at the *ura4* locus under the *fus1* promotor in a *fus1Δ* strain (Figure 1D-F). The control Fus1N-Fus1C was able to support fusion in a manner indistinguishable from the WT (Figure 1E-F). The other chimeric formins instead supported cell fusion to varying reduced degrees (Figure 1E-F). Fus1N-For3C performed very poorly, with only 12% of the cells fused 24h post nitrogen starvation, and exhibited a hazy localization at the region of contact between the two cells. Fus1N-Cdc12C performed relatively well, with a fusion efficiency over 75%, and showed a sharper localization to the region of cell contact. However, in cell pairs that failed to fuse, Fus1N-Cdc12C formed, in the two partner cells, two distinct foci that failed to come together, indicating a position more distant from the membrane than that of WT Fus1 (Movie S1) (Dudin et al., 2015). This unusual localization was due to Cdc12 long C-terminal extension, suggesting it is due to its described function in oligomerisation and actin bundling (Bohnert, Grzegorzewska, et al., 2013), as truncation after Cdc12 FH2 domain led to loss of the double-focus localization and alteration of fusion efficiency (Figure 1E-F). Because neither For3C nor Cdc12C, with or without its C-terminal tail, were able to completely replace Fus1C, these findings suggest a key role of Fus1 actin assembly properties to nucleate the fusion focus. The observation that these chimeras performed better than full-length For3 and Cdc12 also indicates a role of Fus1 N-terminal regulatory region, which we do not address in this work. Here we focus on the specificities contained within the formin C-terminal half.

### Importance of Fus1 expression levels and leucine prototrophy for cell fusion

In the process of repeating the experiments described above with chimeras integrated at the native *fus1* locus, we obtained initially puzzling results. Indeed, these constructs appeared more strongly expressed (Figure 2A, i, compare to Figure 1E) but exhibited a lower fusion efficiency (less than 10% of fused cell pairs 24h post-starvation for both Fus1N-For3C and Fus1N-Cdc12C). Through systematic analysis we resolved that these quantitative discrepancies were accounted for by two variables: the leucine auxotrophy status of the strain and the formin expression level.

**Figure 2.**
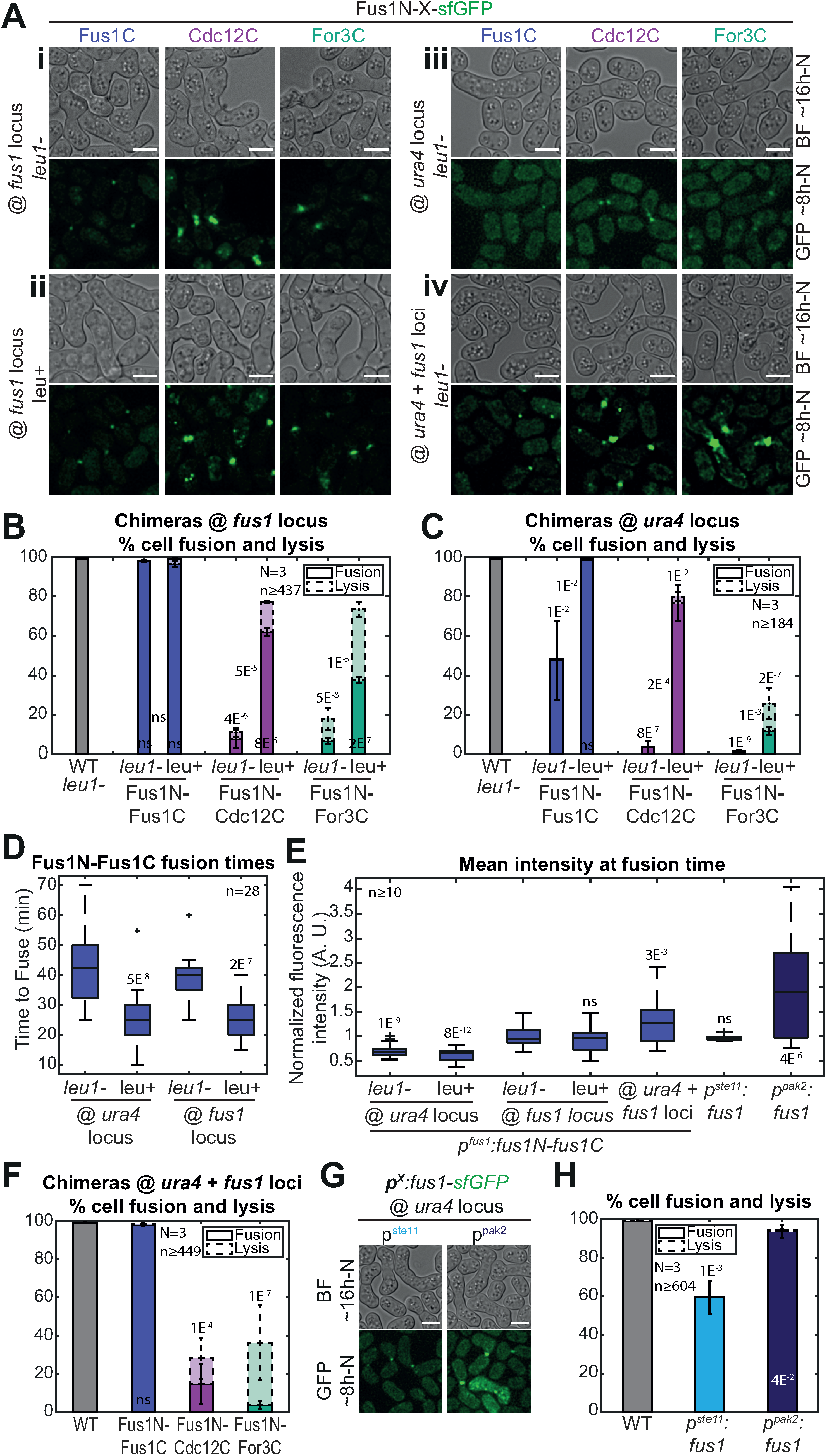
Leucine auxotrophy and formin expression levels have an impact on cell fusion. **A.** DIC images ∼16h post starvation and GFP fluorescence images ∼8h post starvation of *fus1Δ* homothallic strains expressing the formin chimeras described in Figure 1D, either (i) at the *fus1* locus in *leu1-* cells, (ii) at the *fus1* locus in leu+ cells, (iii) at the *ura4* locus in *leu1-* cells, or (iv) at both *ura4* and *fus1* loci in *leu1-* cells. The same chimeras expressed at the *ura4* locus in leu+ cells are shown in Figure 1E. **B.** Percentage of cell pair fusion and lysis 24h after nitrogen removal in strains as in (A, i-ii) compared to the WT. p-values on top of each bar are relative to the WT. p-values between bars compare *leu1-* and leu+ backgrounds. **C.** Percentage of cell pair fusion and lysis 24h after nitrogen removal in strains as in (A, iii) and Figure 1E (same data as Figure 1F shown for comparison here) compared to the WT. p-values on top of each bar are relative to the WT. p-values between bars compare *leu1-* and leu+ backgrounds. **D.** Boxplot of fusion times in strains expressing Fus1N-Fus1C as indicated. The central line indicates the median, the bottom and top edges of the box indicate the 25th and 75th percentiles, respectively, and the whiskers extend to the most extreme data points not considered outliers. **E.** Boxplot of the normalized mean total fluorescence intensity at fusion time in indicated strains. **F.** Percentage of cell pair fusion and lysis 24h after nitrogen removal in strains as in (A, iv) compared to the WT. All strains are *leu1-*. **G.** DIC images ∼16h post starvation and GFP fluorescence images ∼8h post starvation of *fus1Δ* homothallic strains expressing Fus1-sfGFP from the *ste11* or the *pak2* mating-specific promoter at the *ura4* locus. All strains are *leu1-*. **H.** Percentage of cell pair fusion and lysis 24h after nitrogen removal in strains as in (G) compared to the WT. In (B-C, F, H), error bars are standard deviations. All p-values (student’s t-test) are relative to WT unless indicated otherwise. Bars are 5µm.

The major contributor was the leucine status. Indeed, after systematically creating strains with constructs integrated at the *fus1* or the *ura4* locus in a leu+ or *leu1-* background, we found that all *leu1-* strains fused strikingly less well than their leu+ counterparts (Figure 2A-C). Even the Fus1N-Fus1C control exhibited an increase in fusion time in the *leu1-* background compared to its leu+ counterpart, independently of the site of insertion (Figure 2D). While we do not currently understand the reason why leucine auxotrophy is detrimental for cell fusion (uracil and adenine auxotrophies, which are also commonly used in *S. pombe* strains did not show the same effect), these results stress the importance of using strains with identical auxotrophies, which is what we did for this work.

Independently of the auxotrophy effect, we also found that constructs inserted at the *ura4* locus generally exhibited lower fusion efficiencies than constructs inserted at the native *fus1* locus (Figure 2A-C). In fact, the otherwise fully functional Fus1N-Fus1C control was not able to completely support fusion when expressed from the *ura4* locus in a *leu1-* background (Figure 2A,C). Quantification of Fus1N-Fus1C-sfGFP total fluorescence levels at the time of cell fusion, when it is maximal (Dudin et al., 2015), showed a roughly 0.7-fold lower expression from the *ura4* than the *fus1* locus, independently of auxotrophy (Figure 2E). Thus, the genomic context influences the expression of the inserted 1kb-*fus1* promoter, and Fus1 expression levels matter in fusion.

To further probe how formin expression levels influence cell fusion, we constructed a set of strains in the more sensitized *leu1-* background varying Fus1 levels. First, we inserted the Fus1N-forminC constructs at both *ura4* and *fus1* loci (Figure 2A, iv). This yielded a 1.3-fold increased expression, as measured on Fus1N-Fus1C, compared to the *fus1* locus insertion alone (Figure 2E), but had no significant effect on any fusion efficiency (Figure 2A,F). We then expressed full-length Fus1 under *ste11* and *pak2* mating-specific promotors at the *ura4* locus in *fus1Δ* cells (Figure 2G), which respectively gave similar and 2.1-fold higher expression than expression from the endogenous locus (Figure 2E). Unexpectedly, expression from the *ste11* promotor did not allow for a fully successful fusion efficiency, though it was superior to that observed upon lower-level expression under the *fus1* promotor at the same genomic locus (Figure 2H). We hypothesize that cell pairs that fail to fuse express lower levels, and that our measures overestimate the expression level because we only quantified fluorescence in successfully fusing pairs (to ensure quantification was done at the same functional timepoint). Expression from the *pak2* promoter allowed for a much better fusion efficiency, just slightly, but significantly inferior to the endogenous and the dually expressed Fus1 (94%). These results indicate that cell fusion is more robust to increase than reduction in formin levels and suggest that Fus1 native levels are set near the minimum required for function.

To control for these variables, in all subsequent experiments, we systematically used *leu1-* strains with constructs inserted at the native *fus1* locus.

### Fus1 actin assembly properties are tailored to its function

The Fus1N-forminC chimera experiments described above indicate an important role of the formin C-terminal half in achieving cell fusion. The formin C-terminal halves contain FH1 and FH2 domains required for actin assembly, but also a C-terminal regulatory region. In Cdc12, the C-terminal extension is long and contains an oligomerisation domain (Bohnert, Grzegorzewska, et al., 2013). In the formin For3, the C-terminal tail bears the DAD region for auto-inhibition (Martin et al., 2007). The C-terminal regulatory region of Fus1 is short and does not display any detectable DAD domain. To test whether this region of Fus1 is important for cell fusion, we constructed a mutant form lacking the C-terminal regulatory extension, Fus1^ΔCter^. While Fus1^ΔCter^ localized properly to the region of contact between the two cells, it distributed more widely along the contact zone than full-length Fus1 (Figure 3A-C). However, Fus1^ΔCter^ did not cause a significant increase in fusion time (Figure 3D), had only minor effect on fusion efficiency (Figure 3A,E) and showed very mild post-fusion morphogenetic phenotypes (Fig 3A, arrowheads). Because the reduction in fusion efficiency of *fus1^ΔCter^* cells is mild, whereas replacement of the whole Fus1C with either For3C or Cdc12C leads to dramatic loss of function in the same conditions, this suggests that specific aspects of Fus1 FH1 and FH2 domains are critical for cell-cell fusion.

**Figure 3.**
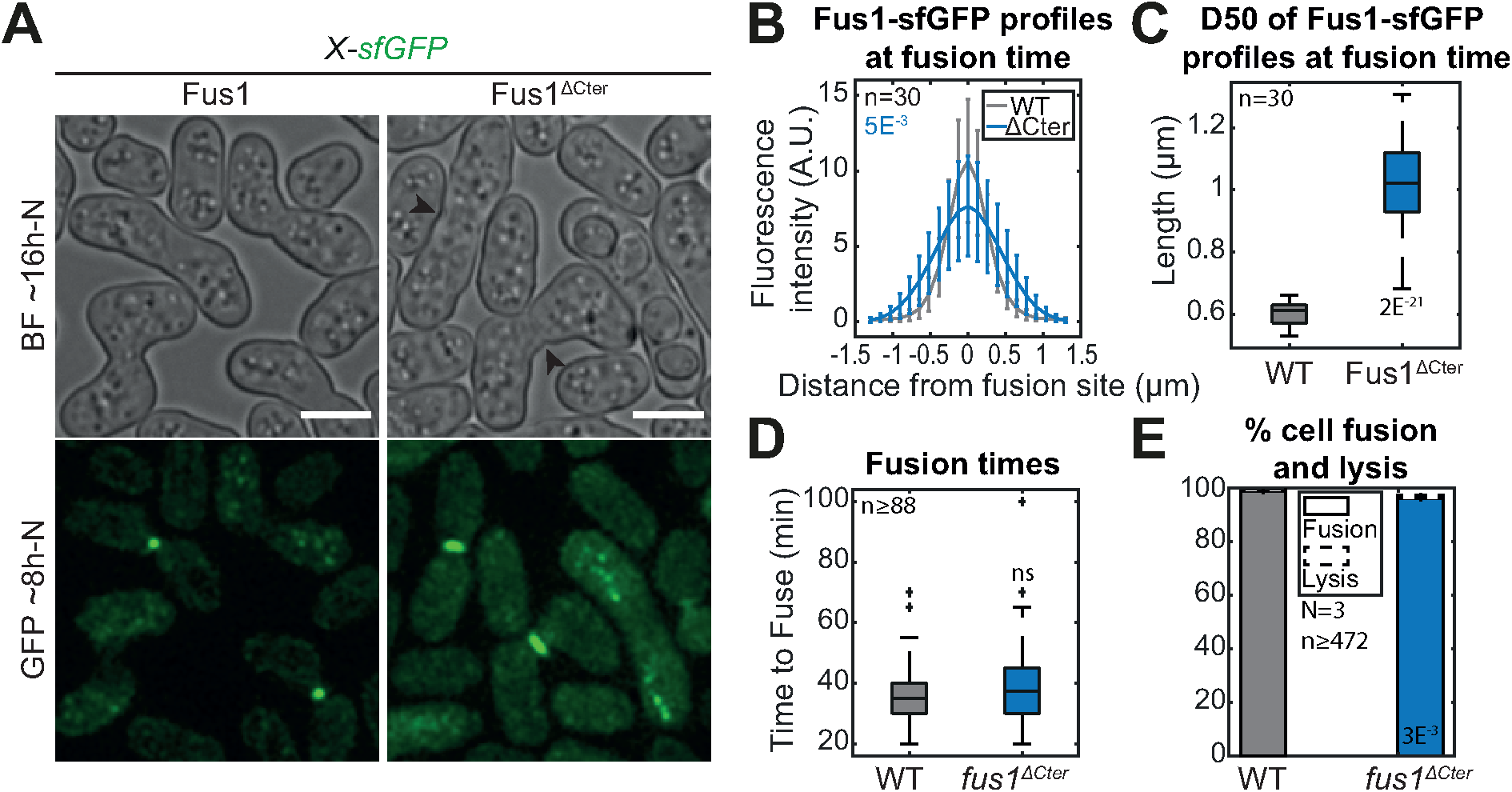
Deletion of Fus1 C-terminal tail leads to spreading of Fus1 localization but has only minor impact on fusion efficiency. **A.** DIC images ∼16h post starvation and GFP fluorescence images ∼8h post starvation of strains expressing sfGFP-tagged Fus1 or Fus1^ΔCter^ from the endogenous *fus1* locus. Fus1^ΔCter^ is cut just after the FH2 domain. Black arrowheads point to cell wall remnants in fused cells. **B.** Profiles of Fus1-sfGFP bleach-corrected fluorescence intensities perpendicular to the cell pair long axis at fusion time in strains as in (A). p-values (student’s t-test) are calculated at the curve maximum. **C.** Boxplot of the width at half maximum of the fluorescence profiles shown in (B). **D.** Boxplot of fusion times in strains as in (A). **E.** Percentage of cell pair fusion and lysis 24h after nitrogen removal in strains as in (A). In (B and E), error bars are standard deviations. All p-values (student’s t-test) are relative to WT. Bars are 5µm.

To examine the specificity in the actin assembly properties of Fus1 FH1-FH2 domains to build the fusion focus, we constructed a set of formin chimeras in which Fus1 N and C-terminal regulatory regions remained constant and only the FH1 and FH2 domains were exchanged between formins. We built upon the extensive *in vitro* characterisation of Fus1, Cdc12 and For3 FH1-FH2 fragments (Scott et al., 2011), which allowed us to control for actin assembly parameters. We first assessed a set of constructs where we kept the FH2 domain of Fus1 and varied the FH1 domain, so as to increase Fus1 elongation speed while keeping the other actin assembly properties constant (Figure 4A). These constructs localized normally to the fusion site (Figure 4B). Doubling the Fus1 FH1 domain or replacing it by half of Cdc12 FH1 domain was previously shown to double actin filament elongation rate, while replacing it by two copies of Cdc12 FH1 was shown to triple it (Scott et al., 2011). This increase in elongation speed was reflected *in vivo* by a larger actin fusion focus, as labelled by Lifeact, which associates with F-actin, or tropomyosin GFP-Cdc8, which associates specifically with formin-assembled actin filaments (Christensen et al., 2017) (Figure 4C), suggesting higher actin filament assembly capacity for the faster formin chimeras. Interestingly, while none of the constructs significantly reduced fusion efficiency at 24h post starvation (Figure 4B,D), they induced a delay in the fusion process. Indeed, the duration of the fusion process, measured from the first formation of the fusion focus to cytosolic mixing, showed an apparent correlation with the formin elongation speed (Figure 4E). We conclude that increasing Fus1 elongation speed is detrimental to its function.

**Figure 4.**
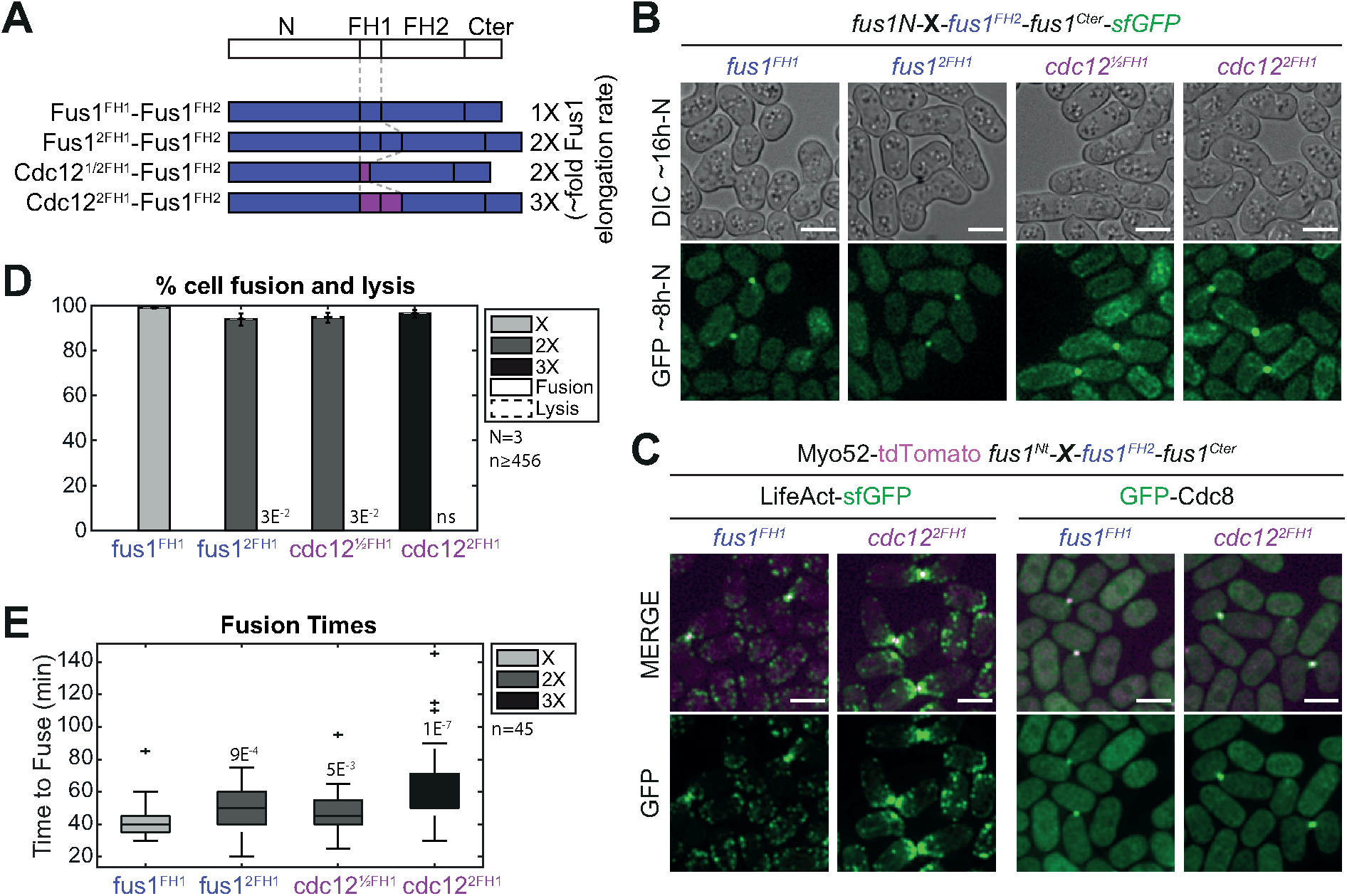
An increase in Fus1 actin elongation rate delays cell fusion. **A.** Scheme of the constructs used in the figure. All constructs were constructed seamlessly (no restriction sites separate domains) and integrated at the endogenous *fus1* locus. As they keep their N- and C-terminal regulatory parts and their FH2 domain constant, they are referred to only by their variable FH1 domain. Indicative actin filament elongation rates as measured *in vitro* on FH1-FH2 fragments by (Scott et al., 2011) are shown on the right, as multiple of Fus1 elongation rate. **B.** DIC images ∼16h post starvation and GFP fluorescence images ∼8h post starvation of homothallic strains expressing formin chimeras as in (A), C-terminally tagged with sfGFP. **C.** Merge and GFP fluorescence images ∼8h post starvation of Myo52-tdTomato (magenta) and either (left) LifeAct-sfGFP or (right) GFP-Cdc8 in homothallic strains expressing untagged formin chimeras with Fus1^FH1^ or Cdc12^2FH1^. **D.** Percentage of cell pair fusion and lysis 24h after nitrogen removal in strains as in (B). Error bars are standard deviations. **E.** Boxplot of fusion times in strains as in (B). All p-values (student’s t-test) are relative to WT. Bars are 5µm.

We then set to examine chimeras with different FH2 domains (Figure 5A-B). Using the Cdc12 FH2 domain and varying FH1 domains, we were able to express in cells formin constructs with a wider range of measured *in vitro* elongation rates than those obtained with Fus1 FH2, including one construct with lower rates than native Fus1. *In vitro* studies showed that Cdc12 has a nucleation rate very close to that of Fus1 (Scott et al., 2011), suggesting that nucleation rate is adequate in these constructs and permitting us to test for an optimum elongation speed in fusion (Figure 5A). The Cdc12^FH1^-Cdc12^FH2^ chimera, with twice the Fus1 elongation speed supports fusion in only 33% of cell pairs (Figure 5B-C). Importantly, reducing the elongation speed to that of native Fus1 by halving Cdc12 FH1 domain led to an important increase in fusion efficiency, with 78% success (Figure 5B-C). By contrast, further decrease (Fus1^FH1^-Cdc12^FH2^) or increase (Cdc12^2FH1^-Cdc12^FH2^) of the elongation rate compromised cell fusion efficiency, to 17 and 15 %, respectively, demonstrating a clear optimum at the native elongation rate of Fus1. The architecture of the fusion focus reflected the elongation rates measured *in vitro*. Indeed, all constructs localized to the site of cell-cell contact (Figure 5B), but the poorest elongator was barely able to form an actin aster, which showed minimal tropomyosin localization (Figure 5D-E). In contrast, the faster elongators formed a F-actin-rich focus from which emanated GFP-Cdc8-decorated actin cables, an organisation not observed with native Fus1 (Figure 5D-E, compare to Figure 4C). These results demonstrate that the native Fus1 elongation rate is best adapted to assemble the fusion focus.

**Figure 5.**
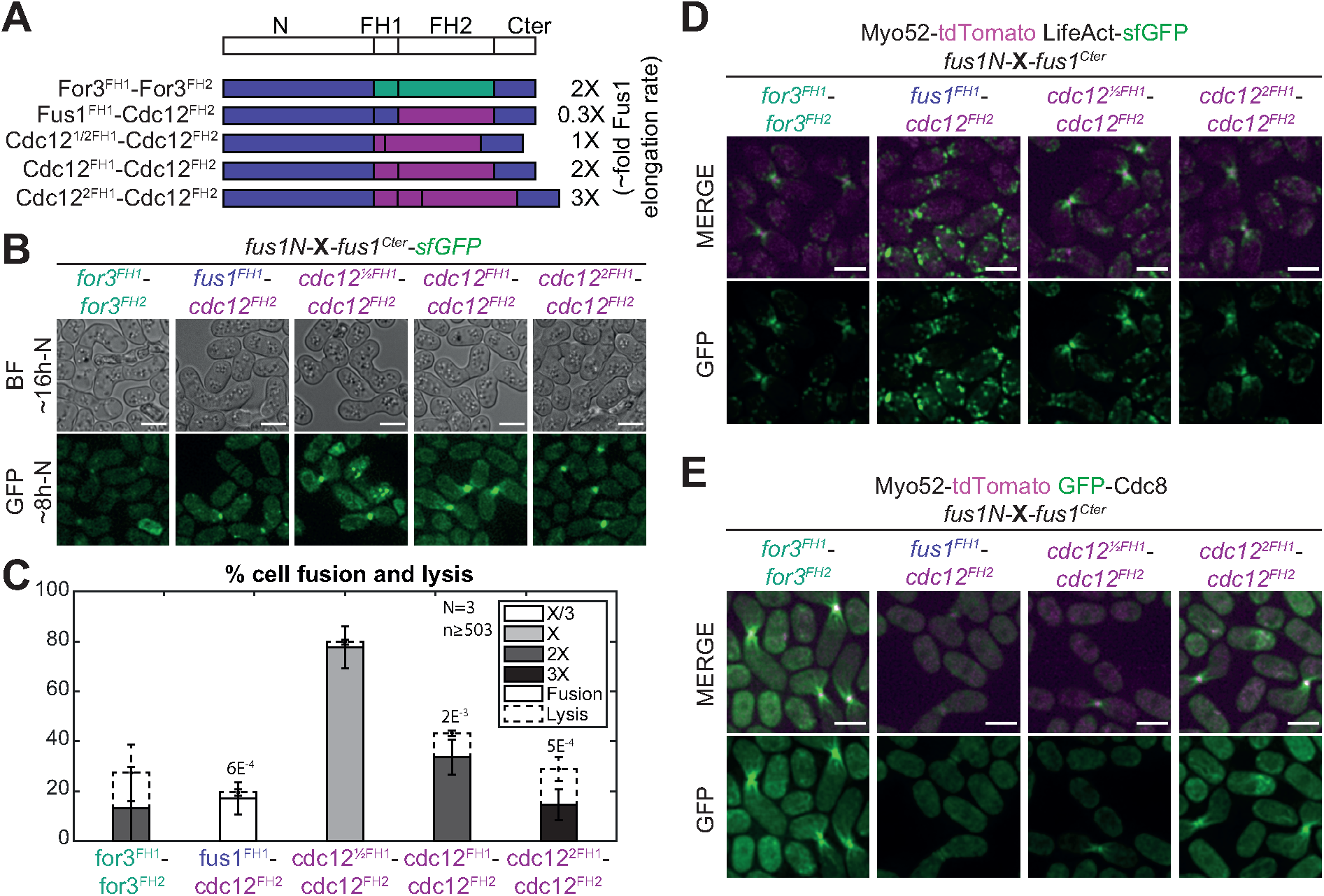
Low elongation and nucleation rates are detrimental for cell fusion and Fus1 contains an additional property within its FH2 domain absent from Cdc12 FH2 domain. **A.** Scheme of the constructs used in the figure. All constructs were constructed seamlessly (no restriction sites separate domains) and integrated at the endogenous *fus1* locus. As they keep their N- and C-terminal regulatory parts constant, they are referred to only by their variable FH1 and FH2 domains. Indicative actin filament elongation rates as measured *in vitro* on FH1-FH2 fragments by (Scott et al., 2011) are shown on the right, as multiple of Fus1 elongation rate. Note that For3 exhibits ∼80-fold lower nucleation rates than Fus1 (not shown). **B.** DIC images ∼16h post starvation and GFP fluorescence images ∼8h post starvation of homothallic strains expressing the formin chimeras shown in (A), C-terminally tagged with sfGFP. **C.** Percentage of cell pair fusion and lysis 24h after nitrogen removal in strains as in (B). Error bars are standard deviations. p-values (student’s t-test) are relative to the most efficient chimera, Cdc12^½FH1^-Cdc12^FH2^. **D.** Merge and GFP fluorescence images ∼8h post starvation of Myo52-tdTomato and LifeAct-sfGFP in homothallic strains expressing untagged formin chimeras shown in (A). **E.** Merge and GFP fluorescence images ∼8h post starvation of Myo52-tdTomato and GFP-Cdc8 in homothallic strains expressing untagged formin chimeras shown in (A). Bars are 5µm.

*In vitro* studies showed that Fus1 is an efficient nucleator (Scott et al., 2011). To test for the importance of actin filament nucleation, we used the For3 FH2 domain, which exhibits a nucleation efficiency over 80-fold lower than Fus1 *in vitro*. A chimera with For3 FH1-FH2 domains (Figure 5A) performed poorly, with barely over 10% of the mating pairs managing to fuse (Figure 5B-C). For3 also elongates actin filaments at about twice the rate of Fus1, and we indeed observed a perturbed actin focus with stronger LifeAct and tropomyosin signal (Figure 5D-E). This stronger signal suggests that the For3 FH1-FH2 chimera either efficiently elongates pre-existing filaments or is a better nucleator *in vivo* than *in vitro*. Some of the inability of For3 FH1-FH2 to support cell fusion may stem from its faster elongation rate. However, compared to the constructs with matched elongation speed described above (Fus1^2FH1^-Fus1^FH2^ and Cdc12^1/2FH1^-Fus1^FH2^ in Figure 4, or Cdc12^FH1^-Cdc12^FH2^ in Figure 5), the poorer performance of the For3 chimera suggests that a high nucleation rate is beneficial for the assembly of the fusion focus.

Together these results show that known changes in the *in vitro* actin assembly properties of the formin Fus1 induce clear alterations in the architecture of the fusion focus *in vivo* and an associated loss of function. This demonstrates that the native actin assembly properties of Fus1 – a relatively low elongation rate (about 5 subunits.s^-1^.µM^-1^) and a relatively high nucleation rate (one filament per two to three dimers) – are tailored to its function in the assembly of the fusion focus.

### Identification of a mutation blocking an additional function in Fus1 FH2 domain

We noted that, even for matching elongation rates, chimeras containing the Fus1 FH2 domain systematically performed better than those containing either For3 or Cdc12 FH2 domain. While for the For3 FH2 chimera this can be rationalized to the importance of efficient nucleation, the comparisons with Cdc12 FH2 chimeras, where both elongation and nucleation rates are matched, suggested to us that Fus1 FH2 contains a specific property critical for cell fusion. The clearest demonstration of this lies in the comparison of Cdc12^2FH1^-Fus1^FH2^ with Cdc12^2FH1^-Cdc12^FH2^, as both constructs differ only through their FH2 domains but perform at 97% and 15%, respectively (Figures 4D and 5C, respectively).

With the aim to identify amino acids required for Fus1 FH2 specific property, we first tested whether it was conserved in Fus1 orthologues. To this aim, we replaced *S. pombe* Fus1 FH2 domain by the corresponding FH2 domains in *Schizosaccharomyces octosporus* (*S. octosporus*) and in the more distant *Schizosaccharomyces japonicus* (*S. japonicus*). We performed this replacement in two distinct construct backgrounds (Figure S1A). First, we used the construct carrying Cdc12^2FH1^, because this is the only FH1 domain to yield identical elongation rates in combination with either Fus1 FH2 or Cdc12 FH2 domain, allowing to control for this variable. This background also results in the most striking difference in cell fusion efficiency between Cdc12 and Fus1 FH2 domains, as noted above, affording better sensitivity to our assay. Second, we used the *S. pombe* WT Fus1 background as a control to ensure that FH2 exchanges (and further mutations introduced below) did not disturb other actin assembly properties. Both the *S. octosporus* and *S. japonicus* FH2 domains were able to replace *S. pombe* FH2 in combination with either Fus1^FH1^ or Cdc12^2FH2^, with localizations and fusion efficiencies equivalent to those conferred by *S. pombe* Fus1 FH2 domain (Figure S1B-C). Thus, the property we are looking for is conserved within the *Schizosaccharomyces* clade.

We used sequence alignments of *Schizosaccharomyces* Fus1 and Cdc12 FH2 domains and homology modelling of *S. pombe* Fus1 and Cdc12 FH2 domains to identify residues conserved in Fus1 but different in Cdc12, and regions with local charge difference (Figures 6A and S1D). The homology models helped eliminate residues likely to affect actin binding, homodimerization or overall formin structure (Figure 6A). Our comparisons identified 2 residues, L^959^ and R^1054^, and 3 poorly conserved regions located in flexible loops, ^935^KEYTG^939^, ^1006^EEVMEV^1011^ and ^1182^NHK^1184^ (numbering and residues refer to *S. pombe* Fus1).

We mutated these residues, replacing them with the corresponding amino acids found in the *S. pombe* Cdc12 sequence, in the Fus1^FH1^ and Cdc12^2FH1^ construct backgrounds described above (Figure 6B). All formins with mutant Fus1 FH2 localized correctly at the site of cell-cell contact (Figure 6C). In the WT background, none of these mutations had any significant impact on fusion efficiency (Figure 6C-D, left), suggesting preservation of dimerization, actin binding and assembly function. In the Cdc12^2FH1^ background, most mutations also had only minor or no effect on fusion efficiencies, but the R1054E mutation in Fus1 FH2 showed strikingly similar phenotypes as the Cdc12 FH2 (Figure 6C-D, right): severely reduced fusion efficiency and a large amount of cell lysis. These cells also showed an altered fusion focus architecture very similar to that caused by *cdc12^2FH1^-cdc12^FH2^*, with cable-like structures originating from the fusion focus (Figure 6E, right), which are absent from the *cdc12^2FH1^-fus1^FH2^* strain (see Figure 4C). The R1054E mutation in Fus1 FH2 also caused the formation of visible, though weaker GFP-Cdc8-labelled cables from the fusion focus in an otherwise WT background (Figure 6E, left). This suggests that the Fus1R1054E FH2 has lost the Fus1-specific property and behaves similarly to Cdc12 FH2 both in terms of fusion focus architecture and fusion/lysis efficiency.

Encouraged by these results, we introduced the complementary mutation in Cdc12 FH2, replacing E^1168^ by the R found in Fus1. This mutation had no marked effect on formin localization or fusion focus architecture of *cdc12^2FH1^-cdc12^FH2^*-expressing cells (Figure S2A-B) but led to a slight rescue of the fusion efficiency (Figure S2A,C), though far lower than WT levels. This is not very surprising and suggests that Cdc12 would require additional mutations – perhaps some of the changes tested above and causing mild phenotypes – to acquire the Fus1-specfic property. Put together, these results suggest that R^1054^ is one of the amino acids involved in the property that makes Fus1 so well equipped to support cell fusion.

### The R1054E mutation does not alter the biochemical properties of Fus1 FH2 on actin *in vitro*

One hypothesis to explain our *in vivo* results is that a particular biochemical property is significantly altered by the R1054E mutation. The reported actin bundling activity of the Fus1 FH2 domain (Scott et al., 2011) appeared as a possible Fus1-specific property not shared with Cdc12 FH2 domain. We therefore expressed and purified Cdc12^2FH1^-Fus1^FH2^ protein constructs that do or do not contain the R1054E mutation (Figure 7A) and tested their actin filament bundling, assembly, and disassembly activities *in vitro* (Figures 7 and S3). Although we were unable to reproduce the bundling activity to the extent described for Fus1 FH1-FH2 with low-speed sedimentation or fluorescence microscopy assays (Scott et al., 2011), both Cdc12^2FH1^-Fus1^FH2^ and Cdc12^2FH1^-Fus1_R1054E_^FH2^ resulted in significantly more pelleted actin than the control actin only samples, indicating moderate bundling activity (Figure 7B-C). However, there was no significant difference in the bundling activity between the two Cdc12^2FH1^-Fus1^FH2^ constructs.

**Figure 6.**
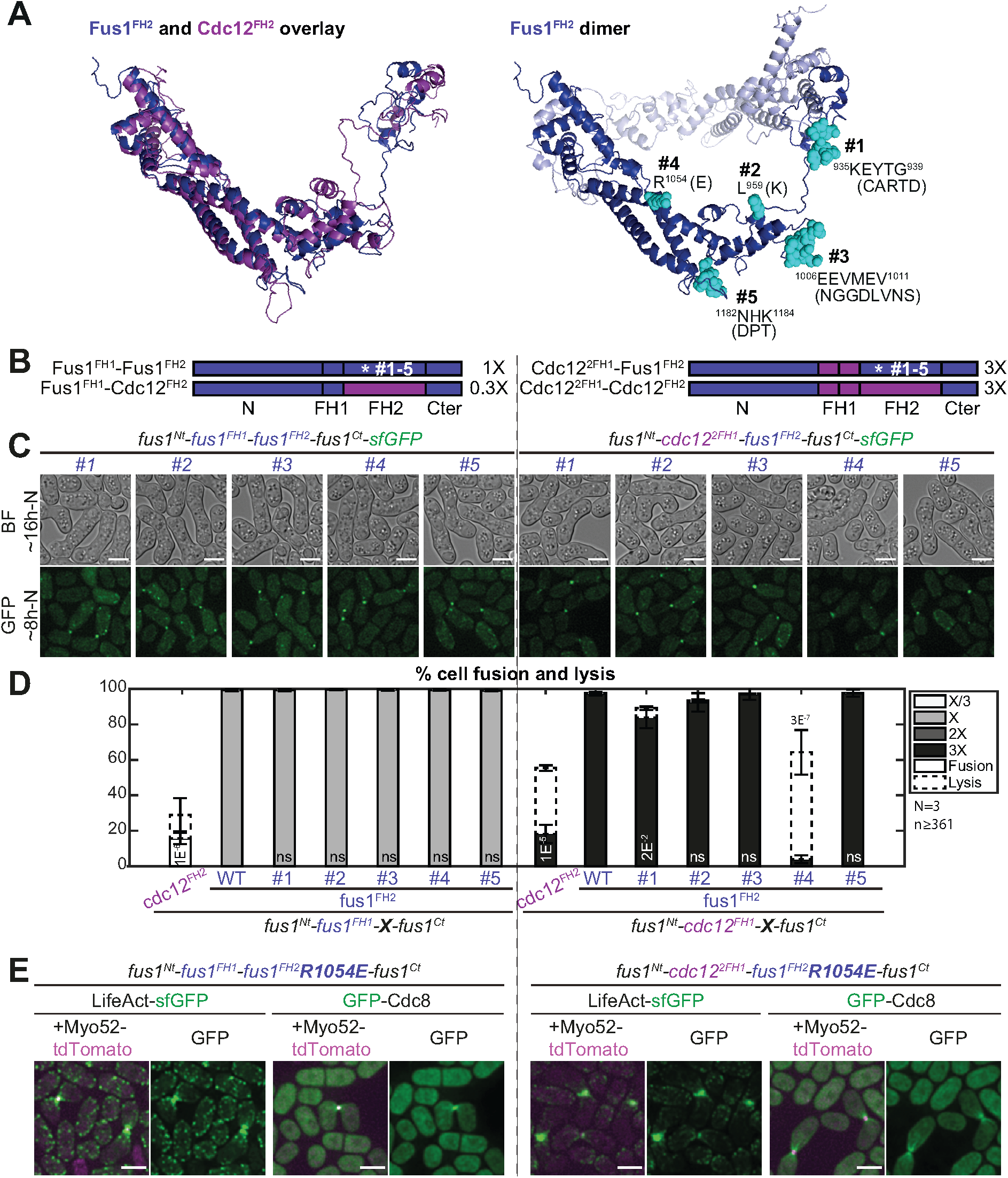
The R1054E mutation in Fus1 FH2 domain partly recapitulates the cell fusion deficiencies observed with Cdc12 FH2. **A.** (Left) Overlay of Fus1 and Cdc12 FH2 domain structures, which were constructed by homology modelling with murine FMNL3. (Right) Dimeric Fus1 FH2 homology model with mutated residues shown in turquoise. Residues were selected in regions that were unlikely to disrupt the formin FH2 dimerization or actin binding, were surface-exposed and had a charge difference between Cdc12 and Fus1 or were in variable loops. **B.** Scheme of the constructs used in the figure. All constructs were constructed seamlessly (no restriction sites separate domains) and integrated at the endogenous fus1 locus. The set of 5 mutations as shown in (A) were introduced in (left) Fus1 or (right) chimeras with Cdc12^2FH1^. The latter shows identical elongation rate when combined with either Fus1^FH2^ or Cdc12^FH2^ (as indicated on the right from measurements *in vitro* on FH1-FH2 fragments by (Scott et al., 2011)). **C.** DIC images ∼16h post starvation and GFP fluorescence images ∼8h post starvation of homothallic strains expressing either (left) mutant Fus1-sfGFP or (right) mutant sfGFP-tagged Cdc12^2FH1^-Fus1^FH2^ formin chimeras. **D.** Percentage of cell pair fusion and lysis 24h after nitrogen removal in strains as in (D) compared to the non-mutated controls. Error bars are standard deviations. p-values (student’s t-test) are relative to the nonmutated Fus1 controls. Chimeras with Cdc12^FH2^ are shown for information but note that Fus1^FH1^-Cdc12^FH2^ (left) cannot be used for direct comparison due to its lower elongation rate. E. Merge and GFP fluorescence images ∼8h post starvation of Myo52-tdTomato and either LifeAct-sfGFP or GFP-Cdc8 in homothallic strains expressing the untagged mutant formins Fus1R1054E (left) or Cdc12^2FH1^-Fus1_R1054E_^FH2^ (right). Note the extended actin network, compared to non-mutated equivalents in Fig 4C and 5E. Bars are 5μm.

**Figure 7.**
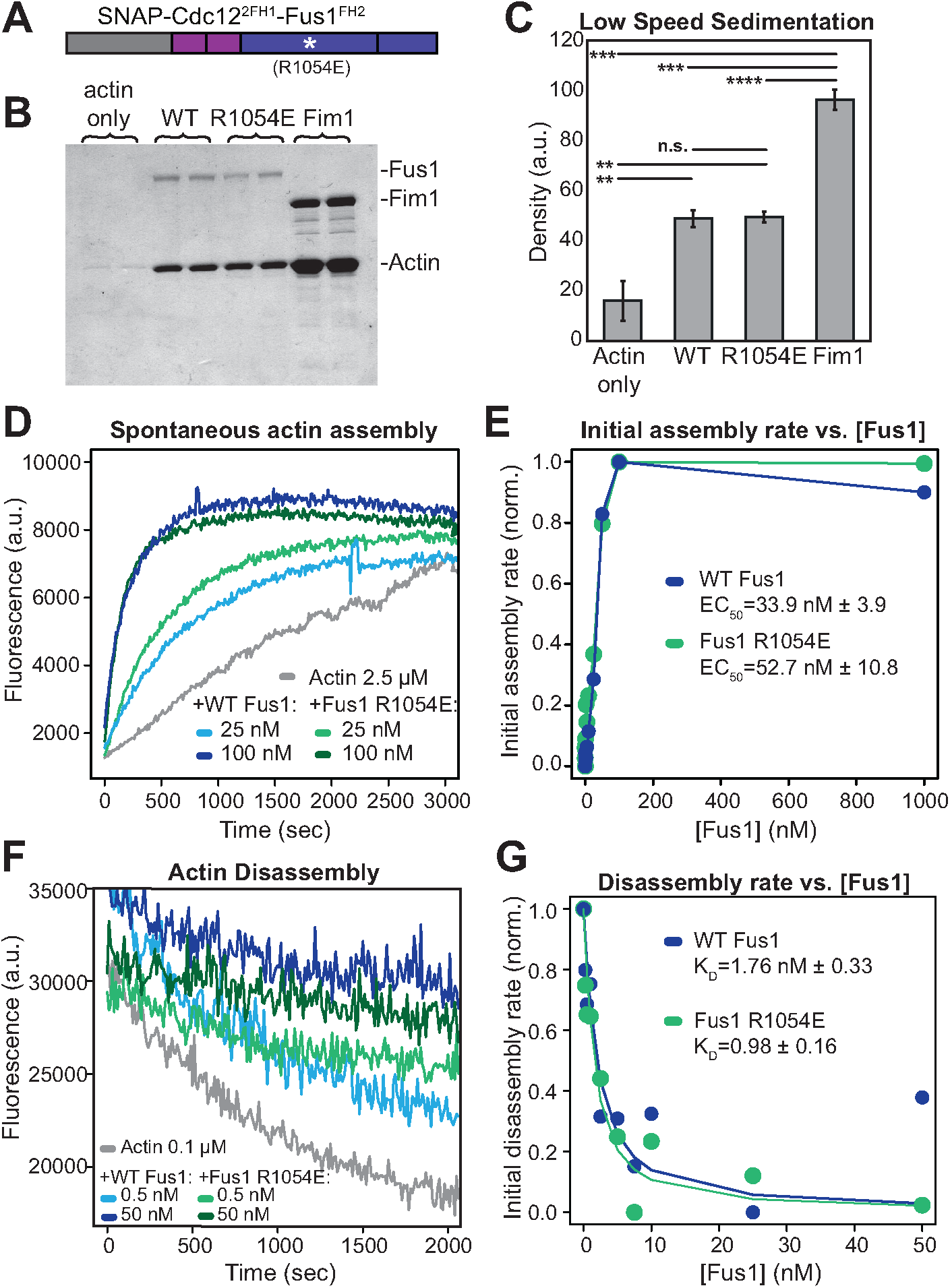
The R1054E mutation does not alter the biochemical properties of Cdc12^2FH1^-Fus1^FH2^ *in vitro*. **A.** Cdc12^2FH1^-Fus1^FH2^ constructs with and without the R1054E mutation utilized for *in vitro* actin biochemistry assays. **B-C.** Low-speed sedimentation of 3 µM preassembled Mg-ATP actin filaments incubated with 1.5 µM of Cdc12^2FH1^-Fus1^FH2^, Cdc12^2FH1^-Fus1_R1054E_^FH2^, or the F-actin bundler Fim1. Samples were spun at 10,000 x g for 20 minutes. **B**. Coomassie Blue-stained gel of pellets. **C**. Graph of the density of actin in the pellets of the assay in (B). Error bars indicate standard errors. ** indicates p < 0.01; *** p < 0.001; **** p < 0.0001 (student’s t-test). **D**-**E**. Spontaneous assembly of 2.5 µM Mg-ATP-actin (10% pyrene-labelled) in the absence and presence of increasing concentrations of Cdc12^2FH1^-Fus1^FH2^ and Cdc12^2FH1^-Fus1_R1054E_^FH2^ . **D.** Representative curves in the absence (grey) and presence of 25 nM formin (light blue and light green) or 100 nM formin (dark blue and dark green). **E.** Representative graph from one of three independent trials of the dependence of the initial assembly slopes (linear phase) on formin concentration. Average effective concentration at half maximal assembly rate (EC50) is reported for each formin ± standard error (p=0.18, n.s., student’s t-test). **F-G**. Barbed end disassembly of preassembled actin filaments (50% pyrene-labelled) upon dilution to 0.1 µM in the absence and presence of increasing concentrations of Cdc12^2FH1^Fus1^FH2^ or Cdc12^2FH1^-Fus1_R1054E_^FH2^ . **F**. Representative curves in the absence (grey) and presence of 0.5 nM formin (light blue and light green) or 50 nM formin (dark blue and dark green). **G**. Representative graph from one of three independent trials of the dependence of the initial disassembly slopes on formin concentration. Average dissociation rate (KD) for the barbed end is reported for each formin ± standard error (p=0.1, n.s., student’s t-test).

We next tested whether these constructs differ in their actin assembly properties by performing spontaneous pyrene actin assembly assays in the presence of varying concentrations of the formin (Figure 7D). By plotting the dependence of the initial actin assembly rates on the concentration of formin, we calculated the concentrations at which the formins achieve half-maximum activity (EC_50_) (Figure 7E). For the wild-type Fus1 chimera, the average EC_50_ from three trials was 33.9 nM ± 3.9, and for the mutant Fus1 chimera, the average EC_50_ from three trials was 52.7 nM ± 10.9 (n.s., p=0.18, student’s t-test) (Figure 7E), indicating that the overall actin assembly properties of these constructs are similar.

To assess the affinity of these formins for actin filament barbed ends, we performed pyrene actin disassembly assays in the presence of varying concentrations of formin (Figure 7F). Pre-assembled actin filaments were diluted to the critical concentration of 0.1 μM, and the rate of barbed end disassembly was followed. Both Cdc12^2FH1^-Fus1^FH2^ and Cdc12^2FH1^-Fus1_R1054E_^FH2^ associated with and reduced disassembly from the barbed end to similar extents (Figure 7F). Furthermore, we plotted the initial disassembly rates against the concentration of the formin, fitted the data using a nonlinear least squares regression, and calculated a dissociation rate (KD), or the affinity of these formins for the barbed end (Figure 7G). We calculated the K_D_ of Cdc12^2FH1^-Fus1^FH2^ to be 1.76 ± 0.33, and the K_D_ of Cdc12^2FH1^-Fus1_R1054E_^FH2^ to be 0.98 ± 0.16 (n.s., p=0.1, student’s t-test), revealing that the wild-type and mutant formin chimeras have similar affinities for actin filament barbed ends.

Lastly, we measured actin filament elongation rates by utilizing TIRF microscopy to directly observe individual filaments assembled by the formins (Figure S3A-B). With both Cdc12^2FH1^-Fus1^FH2^ and Cdc12^2FH1^-Fus1_R1054E_^FH2^, in the absence of profilin two populations of filaments are produced: control filaments (blue arrowheads) that elongate at ∼10 to 13 subunits.s^-1^, and formin-dependent filaments that elongate significantly slower. Cdc12^2FH1^-Fus1^FH2^-associated filaments elongate at 1.1 subunits.s^-1^, consistent with previous results (Scott et al., 2011) (Figure S3B). Cdc12^2FH1^-Fus1_R1054E_^FH2^-associated filaments elongate at 1.0 subunits.s^-1^. In the presence of profilin Cdc3 the elongation rate of control and formin-associated filaments are indistinguishable, but again similar for both Cdc12^2FH1^-Fus1^FH2^ (10.2 subunits.s^-1^) and Cdc12^2FH1^-Fus1^FH2^ (12.8 subunits.s^-1^) (Figure S3B). Furthermore, given that both Cdc12^2FH1^-Fus1^FH2^ and Cdc12^2FH1^-Fus1_R1054E_^FH2^-dependent filaments elongate at nearly identical rates (Figure S3) and have similar ‘bulk’ pyrene actin assembly rates (Figure 7), the actin filament nucleation rates must be similar. These *in vitro* experiments provide strong controls that the reduced functionality of the R1054E mutation *in vivo* is due to changes other than the intrinsic ability to nucleate, elongate, or bundle actin filaments.

## Discussion

How cells can assemble functionally diverse actin structures from a common cytosolic actin pool is a complex question. Actin nucleators confer part of the structure’s identity. Naturally, the different mode of actin nucleation of the Arp2/3 complex and formins yield vastly differing branched and linear networks, but even formin-nucleated actin networks exist in a wide variety of sizes and shapes, which depend on the specific formin nucleator. Regulation of the formin localization and activity is one well-established way to control its time and place of action. Here, we focused on how the formin intrinsic actin assembly properties contribute to the specificity of the assembled actin network architecture. We exploited the simplicity of the fission yeast formin assortment, where only 3 formins each individually assemble a specific actin network, and the prior detailed knowledge on their *in vitro* actin assembly characteristics (Scott et al., 2011), to systematically test the requirements in Fus1 FH1-FH2 domains in assembling the fusion focus. Our results show that 1) Fus1 actin assembly properties are tailored to its function, as reduction or increase in elongation rates, or decrease in nucleation rates are detrimental, and 2) Fus1 FH2 also contains an additional specific property that contributes to conferring the fusion focus its specific architecture.

### Fus1 actin assembly properties are tailored for assembly of the fusion focus

Fus1 exhibits a relatively low elongation rate, at 5 subunits.s^-1^.µM^-1^ *in vitro* (Scott et al., 2011). Our experiments show that this rate is optimal for fusion focus architecture. Indeed, increasing it by providing an FH1 domain with additional polyproline tracks, as in the chimeras with Fus1^FH2^ domain (Figure 4), led to the formation of a larger fusion focus and slowed down the fusion process, proportionally to elongation rate. Similarly, we found that in the chimeras with Cdc12^FH2^ domain (Figure 5), the construct whose elongation rate was matched to that of Fus1 *in vitro* provided the highest fusion efficiency. By contrast, Cdc12^FH2^-based constructs with faster elongation rates formed actin structures with elongated cables and reduced fusion efficiency, again in a manner proportional to elongation rate. These fast-elongating formin chimeras also showed a high percentage of lysed mating pairs. In the context of cells attempting cell fusion, we interpret this as inappropriate cell wall digestion due to loss of spatial precision. The Cdc12^FH2^-based chimeras also allowed us to test the function of a construct with a reduced elongation rate when using the Fus1^FH1^ domain. This slow-elongating formin also did not support function in cell fusion but showed very little cell lysis, likely because it leads to filaments too short to produce a functional actin structure, unable to capture myosin V-associated vesicles for cell wall digestion. Collectively, these experiments demonstrate that Fus1 endogenous elongation rate, which, for a cellular concentration of actin of about 20µM (Wu & Pollard, 2005), corresponds to a filament elongation of about 270 nm.s^-1^, is tailored to its function in fusion focus assembly.

Fus1 is a very potent nucleator, with every other dimer able to initiate a new actin filament *in vitro* (Scott et al., 2011). The observation that For3 FH1-FH2, which has an 80-fold lower nucleation rate *in vitro*, is unable to replace Fus1 and performs worse than Cdc12 with matched elongation rate (compare for instance For3C and Cdc12C constructs in Figure 1, or For3^FH1^-For3^FH2^ and Cdc12^FH1^-Cdc12^FH2^ in Figure 5) suggests that high nucleation rate is beneficial for fusion focus assembly. However, we note that the For3^FH1^-For3^FH2^ chimera efficiently assembled F-actin at the cell-cell contact site, yielding a structure with long actin cables. While we cannot exclude that the nucleation rate may be higher *in vivo* than *in vitro* due to contribution of additional factors – for example, budding yeast nucleation promoting factor Bud6 interacts with the DAD region of formin Bni1 to enhance nucleation by recruiting actin monomers (Graziano et al., 2011) – the For3 chimera may also trigger this actin organisation by elongating pre-existing filaments, for instance at actin patches. The importance of Fus1 nucleation rate should also be viewed in relation with the importance of Fus1 expression levels, reduction of which reduces fusion efficiency, at least in the *leu1-* background. Indeed, the combination of both the nucleation rate and the number of formin molecules will dictate the number of filaments that will assemble at the fusion focus. Together these experiments suggest that high nucleation capacity is required for formation of the fusion focus.

We have not specifically controlled in this study for variations in formin dissociation rate from the filament barbed end, which have also been measured *in vitro*. This is because, whereas the rates were measured on purified simple reactions containing only actin and formins, actin structures are regulated at much faster rates *in vivo* by other actin interacting proteins which either sever the actin filaments or actively unload the formin (Shekhar et al., 2016). Indeed, Fus1 would remain bound to an actin filament for over 25 min and assemble a > 400 µm-long filament according to the dissociation constant measured *in vitro* (Scott et al., 2011), but Fus1-assembled filaments are at most a few µm-long *in vivo*. This difference is due, in particular, to capping protein, which competes with formins for the barbed end of the actin filament, forming a ternary complex which lower the affinity of both proteins for the barbed end (Bombardier et al., 2015; Shekhar et al., 2015), and we have shown competes with Fus1 (Billault-Chaumartin & Martin, 2019).

In the course of our experiments, when initially observing inconsistent phenotypes between strains expressing the same construct, we discovered an implication of leucine auxotrophy in cell fusion, whereby strains prototroph for leucine fuse considerably faster and better than their *leu1-* counterparts. The underlying reason for the effect of leucine auxotrophy is currently unknown and awaits further investigation, but these findings underscore the importance of using identical strain backgrounds, especially during mating, when comparing different strains.

### Fus1 FH2 domain supports an additional function important for cell fusion

The poorer performance of Cdc12 FH2-based constructs, which have similar nucleation rates as Fus1, even with matched elongation rates, suggested to us that the Fus1 FH2 requires an additional, un-attributed property, conserved within the *Schizosaccharomyces* clade, to be fully competent in assembling the fusion focus. The absence of this function in Cdc12 or For3 correlates with the formation of long actin cables emanating from the region of contact between the two cells, especially in fast-elongating constructs. Through sequence alignment, homology modelling and mutational analysis, we were able to identify R1054 in Fus1 as critical for this function. The Fus1 FH2 R1054E mutation has no effect *in vitro* on either elongation or nucleation rates, consistent with the position of this residue on the external face of the FH2 coiled-coil region, not predicted to contact actin. However, it produces phenotypes similar to those observed with Cdc12 or For3 FH2 domains, namely the presence of long cables emanating from the fusion focus. In the context of a fast-elongating construct (with Cdc12^2FH1^), it is also unable to support cell fusion and instead leads to strong cell lysis phenotypes. However, in an otherwise unmodified Fus1, this mutation does not lower fusion efficiency. Collectively, these data suggest that R1054 contributes to the functional specificity of Fus1 FH2, independently of nucleation and elongation rates, but that additional mutations may be necessary to completely abrogate this specific Fus1 role.

One important question is what this role is. We initially thought that bundling activity may be a good candidate. However, the R1054E mutation did not alter the weak bundling activity of the Cdc12^2FH1^-Fus1^FH2^ fragment *in vitro*, indicating that the mutation does not alter the intrinsic bundling activity of Fus1 FH2 domain (Scott et al., 2011). Two arguments support the idea that the mutation may still reduce filament bundling or cross-linking *in vivo*. First, our initial chimeras using the entire C-terminal half of formins (Figure 1) indicate that, in the tested conditions, the C-terminal tail of Cdc12, which contains an oligomerisation domain that promotes actin bundling (Bohnert, Grzegorzewska, et al., 2013), enhances Cdc12 FH1-FH2 function. Second, the extended cables systematically observed for all constructs that did not contain wildtype Fus1 FH2 are consistent with a lack of actin filament cross-linking, yielding filaments able to extend out of the fusion focus nucleation zone. While further work will be required to test this hypothesis, a likely scenario, given the *in vitro* results, is that any bundling activity may be indirect and require additional proteins, potentially binding Fus1 on an interface containing R1054 and absent in our *in vitro* reactions.

The finding that formins are tailored to their cellular function makes intuitive sense in the light of evolution, where selection of the most adapted parameters may have been refined through generations. However, a recent review concluding with open questions on formins stated that “perhaps most challenging of all: how does possessing particular actin polymerization activities render a formin isoform most suitable to fulfil its cellular role?” (Courtemanche, 2018). Our work, along with other recent work on *S. pombe* Cdc12 (Homa et al., 2021) and *Physcomitrella patens* formin For2 (Vidali et al., 2009), contributes to establishing how the formin actin assembly activities are customised to build specific, functional actin structures.

## Materials and methods

### Strain construction

Strains were constructed using standard genetic manipulation of *S. pombe* either by tetrad dissection or transformation and can be found in table S1. Oligonucleotides and plasmids used can be found in tables S2 and S3, respectively, with details on how the plasmids were constructed.

myo52-tdTomato:natMX and fus1-sfGFP:kanMX tags were constructed by PCR-based gene targeting of a fragment from a template pFA6a plasmid containing the appropriate tag and resistance cassette, amplified with primers carrying 5’ extensions corresponding to the last 78 coding nucleotides of the ORF and the first 78 nucleotides of the 3’UTR, which was transformed and integrated in the genome by homologous recombination, as previously described (Bähler et al., 1998). Similarly, fus1Δ::hphMX was constructed by PCR-based gene targeting of a fragment from a template pFA6a plasmid containing the appropriate resistance cassette, amplified with primers carrying 5’ extensions corresponding to the last 78 nucleotides of the 5’UTR and the first 78 nucleotides of the 3’UTR, which was transformed and integrated in the genome by homologous recombination. fus1^Δ1278-1372^-sfGFP:kanMX was constructed by PCR-based gene targeting of a fragment from a template pFA6a plasmid with primers carrying 5’ extensions corresponding to the 78 nucleotides upstream of the deleted region and the first 78 nucleotides of the 3’UTR, which was transformed and integrated in the genome by homologous recombination.

Construction of the strains expressing formin constructs from the *fus1* promotor at the *ura4* locus as a multicopy integration (ura4-294:p^fus1^:fus1-sfGFP:ura4+, ura4-294:p^fus1^:cdc12-sfGFP:ura4+, ura4-294:p^fus1^:for3-sfGFP:ura4+ in Figure 1A-C) was done by homologous recombination of a transformed ura4^EndORF^-ura4^3’UTR^-p^fus1^-ForminConstruct-sfGFP-ura4^StartORF^-ura4^5’UTR^ fragment, obtained from StuI digestion of a pRIP based plasmid (pSM1656, pSM1658, pSM1657). Such recombination recreates a new integration site, which has been shown to be unstable and can lead to multiple insertion (Vještica et al., 2020), which is why we switched to single integration vectors for the rest of the study.

Construction of the strains expressing Fus1N-Fus1C, Fus1N-Cdc12C, Fus1N-For3C and Fus1N-Cdc12CΔC chimeras from the *fus1* promotor at the *ura4* locus as a single integration (ura4+:p^fus1^:fus1N^1-792^-fus1C^793-1372^-sfGFP, ura4+:p^fus1^:fus1N^1-792^-cdc12C^888-1841^-sfGFP, ura4+:p^fus1^:fus1N^1-792^-for3C^715-1461^-sfGFP, ura4+:p^fus1^:fus1N^1-792^-cdc12CΔolig^888-1451^-sfGFP) was done by homologous recombination of a transformed ura4^5’UTR^-ura4^ORF^-ura4^3’UTR^-p^fus1^-ForminConstruct-sfGFP-ura4^3’’^ fragment, obtained from PmeI digestion of a pUra4^PmeI^ based plasmid (pSM2594, pSM2595, pSM2604 and pSM2596, respectively). This leads to a stable single integration at the *ura4* locus (Vještica et al., 2020).

Construction of the strains expressing fus1 under either *ste11* or *pak2* promotor at the *ura4* locus as a single integration (ura4+:p^ste11^:fus1-sfGFP and ura4+:p^pak2^:fus1-sfGFP) was done by homologous recombination of a transformed ura4^5’UTR^-ura4^ORF^-ura4^3’UTR^-p^ste11 or pak2^-Fus1-sfGFP-ura4^3’’^ fragment, obtained from PmeI digestion of a pUra4^PmeI^ based plasmid (pSM2828 and pSM2829, respectively).

Construction of the strains expressing tagged formin constructs from the endogenous locus (fus1N^1-792^-fus1C^793-1372^-sfGFP:kanMX, fus1N^1-792^-cdc12C^888-1841^-sfGFP:kanMX, fus1N^1-792^-for3C^715-1461^-sfGFP:kanMX, fus1^1-868^-fus1^792-1372^-sfGFP:kanMX, fus1^1-791^-cdc12^928-972^-fus1^869-1372^-sfGFP:kanMX, fus1^1-791^-cdc12^882-972^-cdc12^882-972^-fus1^869-1372^-sfGFP:kanMX, fus1^1-791^-for3^718-1265^-fus1^1278-1372^-sfGFP:kanMX, fus1^1-868^-cdc12^973-1390^-fus1^1278-1372^-sfGFP:kanMX, fus1^1-791^-cdc12^928-1390^-fus1^1278-1372^-sfGFP:kanMX, fus1^1-791^-cdc12^882-1390^-fus1^1278-1372^-sfGFP:kanMX, fus1^1-791^-cdc12^882-972^-cdc12^882-1390^-fus1^1278-1372^-sfGFP:kanMX, fus1^1-868^-Sjfus1^908-1317^-fus1^1278-1372^-sfGFP:kanMX, fus1^1-868^-Sofus1^857-1265^-fus1^1278-1372^-sfGFP:kanMX, fus1^1-792^-cdc12^882-972^-cdc12^882-972^-Sjfus1^908-1317^-fus1^1278-1372^-sfGFP:kanMX, fus1^1-792^-cdc12^882-972^-cdc12^882-972^-Sofus1^857-1265^-fus1^1278-1372^-sfGFP:kanMX, fus1(KEYTG935-939CARTD)-sfGFP:kanMX, fus1(L959K)-sfGFP:kanMX, fus1(EEVMEV1006-1011NGGDLVNS)-sfGFP:kanMX, fus1(R1054E)-sfGFP:kanMX, fus1(NHK1182-1184DPT)-sfGFP:kanMX, fus1^1-792^-cdc12^882-972^-cdc12^882-972^-fus1^869-1372^(KEYTG935-939CARTD)-sfGFP:kanMX, fus1^1-792^-cdc12^882-972^-cdc12^882-972^-fus1^869-1372^(L959K)-sfGFP:kanMX, fus1^1-792^-cdc12^882-972^-cdc12^882-972^-fus1^869-1372^(EEVMEV1006-1011NGGDLVNS)-sfGFP:kanMX, fus1^1-792^-cdc12^882-972^-cdc12^882-972^-fus1^869-1372^(R1054E)-sfGFP:kanMX, fus1^1-792^-cdc12^882-972^-cdc12^882-972^-fus1^869-1372^(NHK1182-1184DPT)-sfGFP:kanMX, fus1^1-792^-cdc12^882-972^-cdc12^882-1390^(E1168R)-fus1^1278-1372^-sfGFP:kanMX) was done by homologous recombination of a transformed fus1^5’UTR^-ForminConstruct-sfGFP-kanMX-fus1^3’UTR^ fragment, obtained from a gel purified, SalI and SacII or AatII and SacII (Cdc12 FH2 contains an endogenous SalI site) digested pFA6a based plasmid (pSM2690, pSM2691, pSM2692, pSM2621, pSM2702, pSM2700, pSM2631, pSM2701, pSM2632, pSM2622, pSM2699, pSM2855, pSM2857, pSM2854, pSM2856, pSM2849, pSM2850, pSM2851, pSM2852, pSM2853, pSM2844, pSM2845, pSM2846, pSM2847, pSM2848 and pSM2924, respectively).

Similarly, construction of the strains expressing untagged formin constructs from the endogenous locus (fus1:kanMX, fus1^1-791^-cdc12^882-972^-cdc12^882-972^-fus1^869-1372^:kanMX, fus1^1-791^-for3^718-1265^-fus1^1278-1372^:kanMX, fus1^1-868^-cdc12^973-1390^-fus1^1278-1372^:kanMX, fus1^1-791^-cdc12^928-1390^-fus1^1278-1372^:kanMX, fus1^1-792^-cdc12^882-972^-cdc12^882-1390^-fus1^1278-1372^:kanMX, fus1(R1054E):kanMX, fus1^1-792^-cdc12^882-972^-cdc12^882-972^-fus1^869-1372^(R1054E):kanMX, fus1^1-792^-cdc12^882-972^-cdc12^882-1390^(E1168R-fus1^1278-1372^):kanMX) was done by homologous recombination of a transformed fus1^5’UTR^-ForminConstruct-sfGFP-kanMX-fus1^3’UTR^ fragment, obtained from a gel purified, SalI and SacII or AatII and SacII digested pFA6a based plasmid (pSM2913, pSM2914, pSM2918, pSM2915, pSM2916, pSM2917, pSM2961, pSM2962 and pSM2963, respectively).

leu1-32:p^nmt41^:GFP-cdc8:ura4+ (Skoumpla et al., 2007), fus1Δ::LEU2+ (Petersen et al., 1998b) and ade6+:p^act1^:LifeAct-sfGFP:term^ScADH1^:bsdMX (Vještica et al., 2020) trace back to the aforementioned papers and are kind gifts from the afore mentioned labs.

### Growth conditions

For mating experiments, homothallic (*h90*) strains able to switch mating types were used, where cells were grown in liquid or agar Minimum Sporulation Media (MSL), with or without nitrogen (+/-N) (Egel et al., 1994; Vjestica et al., 2016).

Live imaging of *S. pombe* mating cells protocol was adapted from (Vjestica et al., 2016). Briefly, cells were first pre-cultured overnight in MSL+N at 25°C, then diluted to OD600 = 0.05 into MSL+N at 25°C for 20 hours. Exponentially growing cells were then pelleted, washed in MSL-N by 3 rounds of centrifugation, and resuspended in MSL-N to an OD600 of 1.5. Cells were then grown 3 hours at 30°C to allow mating in liquid, added on 2% agarose MSL-N pads, and sealed with VALAP. We allowed the pads to rest for 30 min at 30°C before overnight imaging or for 21h at 25°C for 24h post-starvation fusion efficiencies snapshot imaging, respectively.

### Live imaging microscopy

Images presented in Figures 1, 2, 3, 4, 5, 6, S1 and S2 were obtained using a DeltaVision platform (Applied Precision) composed of a customized inverted microscope (IX-71; Olympus), a UPlan Apochromat 100×/1.4 NA oil objective, a camera (CoolSNAP HQ2; Photometrics or 4.2Mpx PrimeBSI sCMOS camera; Photometrics), and a color combined unit illuminator (Insight SSI 7; Social Science Insights). Images were acquired using softWoRx v4.1.2 software (Applied Precision). Images were acquired every 5 minutes during 9 to 15 hours. To limit photobleaching, overnight videos were captured by optical axis integration (OAI) imaging of a 4.6-μm z-section, which is essentially a real-time z-sweep.

### Quantification and statistical analysis of live imaging data

Percentages of cell fusion and lysis as in Figures 1C, 1F, 2B, 2C, 2F, 2H, 3E, 4D, 5C, 6D, S1C and S2C were calculated as in (Dudin et al., 2015). Briefly, 24h post-starvation, fused cell pairs, lysed pairs and the total number of cell pairs were quantified using the ImageJ Plugin ObjectJ, and percentages were calculated using the following equations:

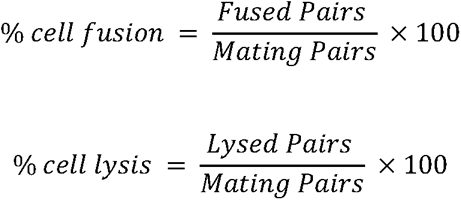

Fusion Times as in Figures 2D, 3D and 4E were calculated in overnight time lapse movies at 5-minutes interval using the 2-dot Myo52-tdTomato stage (Dudin et al., 2015) as a marker for the beginning of the fusion process and either the entry of GFP expressed under control of the P-cell-specific *p^map3^* promoter into the *h-* partner, or the maximum intensity of the Myo52-tdTomato dot, the two of which perfectly correlate (Dudin et al., 2015), as a marker for the end of the process.

Fusion Focus intensities at fusion time as in Figure 3B were obtained in overnight time lapse movies at 5-minutes interval using either the entry of GFP into the *h-* partner, or the maximum intensity of the Myo52-tdTomato dot to determine the moment of fusion. At that time frame, a fluorescence profile across the fusion focus perpendicular to the long axis of the mating pair was recorded. Profiles were background-subtracted and corrected for bleaching as follows: First, the cell fluorescence intensity was recorded over time in a square of about 7x7 pixels in 12 control (non-mating) cells. These fluorescence profiles were averaged, and the mean was fitted to a double exponential as it was describing our data better (Vicente et al., 2007):

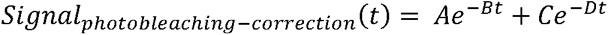

We then used this fit to correct the fluorescence profiles across the fusion focus for photobleaching. After subtracting background signal, the value at each timepoint was divided by the photo-bleaching correction signal:

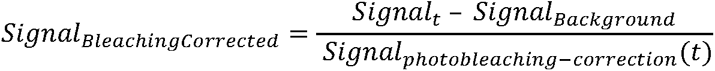

Corrected profiles were then averaged and plotted. Widths at Half maximum (D50) as in Figure 3C were then calculated using these fluorescence profiles. This was done for each cell and then plotted as a boxplot.

Total fluorescence intensities as in Figures 2E in mating pairs at fusion time were obtained using 5-minute time lapse overnight movies where fusion time was assessed for each mating cell, by outlining the mating pairs and recoding the mean fluorescence intensity for each of them at the determined time point. The signal was bleaching-corrected as described above.

All plots, fittings, corrections and normalisations in Figures 1, 2, 3, 4, 5, 6, S1 and S2 were made using MATLAB home-made scripts. For boxplots, the central line indicates the median, and the bottom and top edges of the box indicate the 25th and 75th percentiles, respectively. The whiskers extend to the most extreme data points not considering outliers. For bar plots, error bars represent the standard deviation. Statistical p-values were obtained using a two-sided t-test, after normal distribution had been visually checked using a simple histogram. No further verification was made to ascertain that the data met assumptions of the statistical approach. All values below 0.05 are mentioned in the figures, including sample size.

### Homology modelling

The suitable templates for Fus1 and Cdc12 structure modelling in Figure 6A were found using the HHpred tool (Zimmermann et al., 2018). The models of Fus1 protein were calculated based on the template of diaphanous protein from mice, stored under the 3OBV code in the Protein Data Bank (Otomo et al., 2010). The sequence identity between these protein is 20%. The yeast S. cerevisiae Bni1 protein structure, stored under the 5UX1 code in Protein Data Bank, served as a template for Cdc12 protein (Xu et al., 2004). Both sequences share 34% of sequence identity. For the modelling of Fus1 and Cdc12 protein dimers we used the dimeric structure of FMNL3 protein bound to actin, stored in the PDB under the 4EAH code (Thompson et al., 2013). Fus1 shared around 23% and Cdc12 around 25% of sequence identity with this template, respectively. The models’ structures were calculated using Modeller 9v18 program (Sali & Blundell, 1993). The models of Fus1 and Cdc12 proteins were aligned with UCSF Chimera visualization program (Pettersen et al., 2004). The alignments points, where opposite charge residues were present in both sequences or where a group of charged amino acid appeared in only one of the sequences, were identified. We took into account the fragments that were exposed on the surface of the models - and consequently accessible for interactions with other proteins - and not interfering with dimerization of the domain or with actin elongation.

### Protein expression and purification

Actin was purified from rabbit skeletal-muscle acetone powder (Spudich & Watt, 1971) and labelled on Cys374 with pyrenyliodoacetamide (Pollard & Cooper, 1984), or lysines with Alexa488-succinimidylester (Isambert et al., 1995). Fission yeast profilin Cdc3 was overexpressed and purified from Escherichia coli (Lu & Pollard, 2001). Fimbrin Fim1 was expressed in Escherichia coli and purified via His-tag affinity to Talon Metal Affinity Resin (Clontech, Mountain View, CA) (Skau & Kovar, 2010). SNAP-tagged Cdc12^2FH1^-Fus1^FH2^ and Cdc12^2FH1^-Fus1_R1054E_^FH2^ constructs (including the C-terminal tail) were expressed in Escherichia coli strain BL21-Codon Plus (DE3) RP (Agilent Technologies, Santa Clara, CA) with 0.5 mM isopropyl β-D-1-thiogalactopyranoside for 16 h at 16 °C. Cells were lysed with sonication in extraction buffer [50 mM NaH2PO4 (anhydrous), 500 mM NaCl, 10% glycerol, 10 mM imidazole, 10 mM BME, pH 8] with EDTA-free Protease Inhibitor Cocktail (Roche, Basel, Switzerland) and were clarified by centrifugation. The extract was incubated for 1 h at 4 °C with Talon resin (Clontech, Mountain View, CA), loaded onto a column, washed with extraction buffer, and the protein was eluted with 250 mM imidazole. Formin proteins were dialyzed into SNAP buffer [20 mM HEPES (pH 7.4), 200 mM KCl, 0.01% NaN3, 10% glycerol, and 1 mM DTT] and filtered on a Superdex 200 10/300 GL column (GE Healthcare, Little Chalfont, UK). Aliquots of the protein were flash-frozen in liquid nitrogen and stored at -80 °C.

### Low-speed sedimentation

F-actin bundling activity was determined from low-speed sedimentation assays. Mg-ATP-actin (15 μM) was preassembled for 1 hr at 25°C, then 3.0 μM was aliquoted into Eppendorf tubes and incubated with 1.5 μM of Cdc12^2FH1^-Fus1^FH2^, Cdc12^2FH1^-Fus1_R1054E_^FH2^, or fission yeast fimbrin (Fim1) for 20 min at 25°C. Samples were spun at 10,000 × *g* for 20 min at 25°C. Pellets were resuspended in sample buffer and separated by 12.5% SDS–PAGE and stained with Coomassie Blue (Figure 7B). The density of protein bands as in Figure 7C was analysed with ImageJ.

### Pyrene assembly and disassembly assays

Assembly and disassembly of actin filaments were measured by observing changes in pyrene fluorescence over time. Fluorescence of 10%-labelled pyrene actin (excitation 364 nm, emission 407 nm) was measured with m200Pro (Tecan) fluorescent plate reader. Final protein concentrations are indicated in the figure legends. For spontaneous assembly assays, 15 μM 10% pyrene-labeled Mg-ATP-actin with 100x anti-foam 204 (0.005%; Sigma) was added to the upper row of a 96-well nonbinding black plate (Corning, Corning, NY). A range of concentrations of formin, plus 10X KMEI [500 mM KCl, 10 mM MgCl_2_, 10mM EGTA, 100 mM imidazole (pH 7.0)] and Mg-buffer G [2 mM Tris (pH 8.0), 0.2 mM ATP, 0.1 mM MgCl_2_, 1 mM NaN_3_, 0.5 mM DTT] was added to the lower row of the plate. Reactions were initiated by transferring the contents of the lower wells to the upper wells with a 12-channel pipet, and fluorescence was read in the upper wells.

For depolymerization assays, a 5.0 μM mixture of 50% pyrene-labeled Mg-ATP-actin monomers was preassembled for 1 hr in the upper row of a 96-well nonbinding black plate. Protein, 10X KMEI, and SNAP buffer [20 mM HEPES (pH 7.4), 200 mM KCl, 0.01% NaN3, 10% glycerol, and 1 mM DTT] were placed in a lower row of the plate. Reactions were initiated by mixing the contents of the lower wells with the pre-polymerized filaments, which diluted the actin to 0.1 μM.

### Analysis of pyrene data

Fluorescence data as in Figure 7D and 7F was plotted in Rstudio (https://www.rstudio.com/). Initial assembly rates as in Figure 7E were calculated by finding the slope of a linear regression to the linear portions of the graph from 0-150 seconds. Initial depolymerization rates as in Figure 7G were calculated by finding the slope of a linear regression to the linear portions of the graphs from 0-500 seconds.

Dissociation rates (K_D_) were calculated by fitting the depolymerization data using the nonlinear least squares function, and solving the equation:

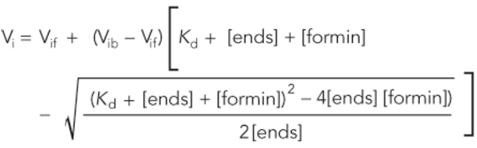

V_i_ is the observed elongation or depolymerization rate, V_if_ is the elongation or depolymerization rate when barbed ends are free, V_ib_ is the elongation or depolymerization rate when barbed ends are bound, [ends] and [formin] are barbed-end and formin concentrations.

### TIRF microscopy

Mg-ATP-Actin (10% Alexa 488-labeled) polymerization was triggered in presence of 10 mM imidazole (pH 7.0), 50 mM KCl, 1 mM MgCl2, 1 mM EGTA, 50 mM DTT, 0.2 mM ATP, 50 mM CaCl2, 15 mM glucose, 20 mg/mL catalase, 100 mg/mL glucose oxidase, and 0.5% (400 centipoise) methylcellulose. This polymerization mix was complemented with proteins of interest as indicated in the figure legends. TIRF-microscopy images as in Figure S3A were acquired using an Olympus IX-71 microscope through TIRF illumination, recorded with a iXon EMCCD camera (Andor Technology), and a cellTIRF 4Line system (Olympus). Actin filament elongation rates as in Figure S3B were measured using the ImageJ software (NIH, Bethesda, Maryland, USA, http://imagej.nih.gov/ij/, 1997-2015). To compare Cdc12^2FH1^-Fus1^FH2^ elongation rates to previously reported rates (Scott et al., 2011), we used the same normalization method where rates were adjusted based on normalization of internal control filaments to 10.0 subunits.s^−1^.μM^−1^.

## Acknowledgements

We thank Daniel Mulvihill, Iain Hagan and Aleksander Vještica for sharing strains as well as Boris Sieber for careful reading of the manuscript. This work was funded by grants from the Swiss National Science Foundation (310030B_176396 and 310030_191990) and the European Research Council (CoG CellFusion) to SGM, National Institutes of Health’s Molecular and Cellular Biology Training Grant T32 GM007183 to SY, and National Institutes of Health’s Grant R01 GM079265 to DRK.

## Author contributions

IBC and SGM conceived the project. SGM performed the experiments in Figure 1B. CAA, SEY and CS performed the experiments in Figures 7 and S3. JI and VZ performed the molecular modelling part of the project. IBC performed all other experiments with technical assistance from LM. SGM and DRK acquired funding. SGM coordinated the project. IBC and SGM wrote the first draft of the manuscript, which was revised by all authors.

---

**Figure S1 (related to Figure 6).**
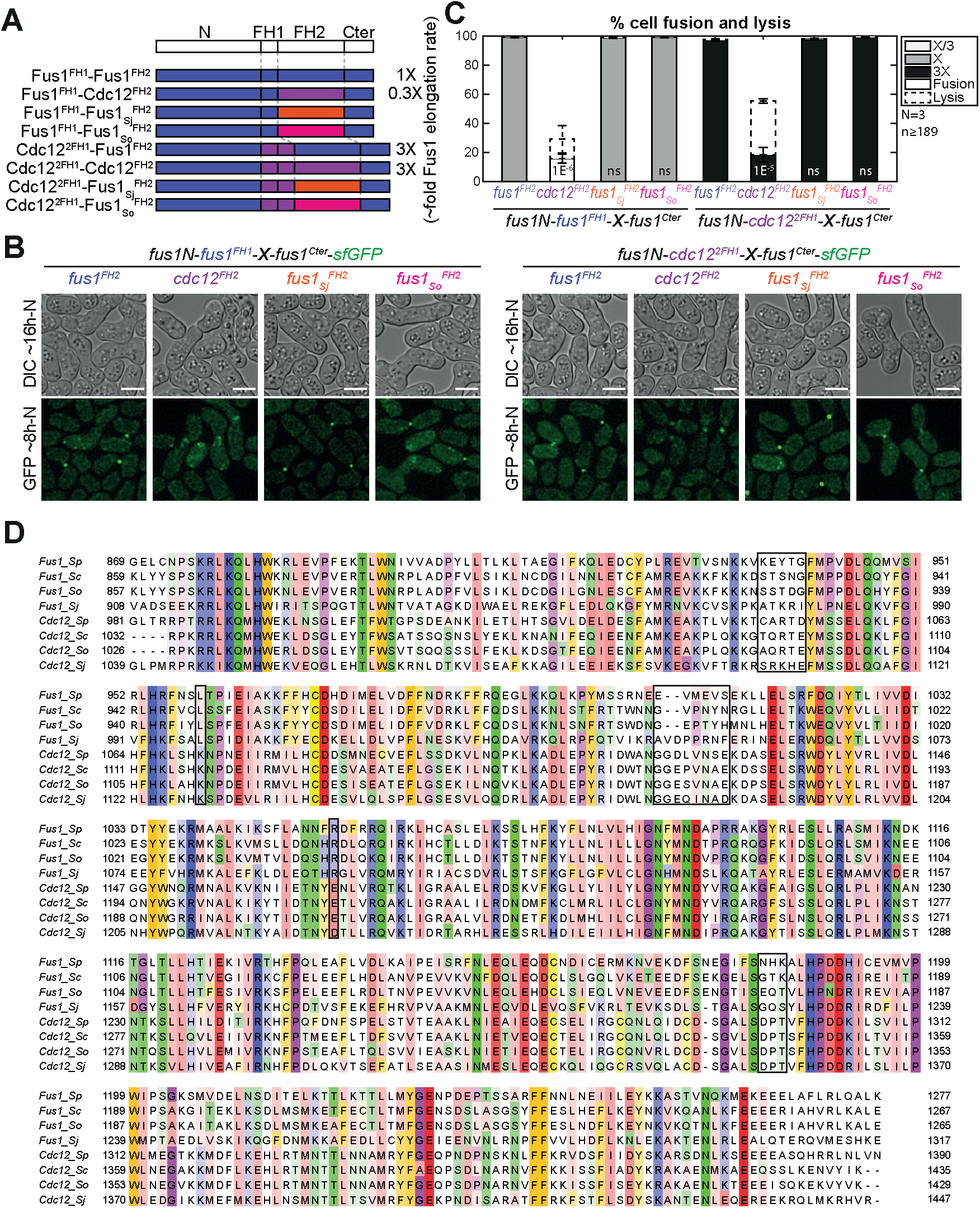
Fus1 FH2 specific property is conserved within the *Schizosaccharomyces* clade. **A.** Scheme of the constructs used in the figure. All constructs were constructed seamlessly (no restriction sites separate domains) and integrated at the endogenous *fus1* locus. As they keep their N- and C-terminal regulatory parts constant, they are referred to only by their FH1 and FH2 domains. Where known, indicative actin filament elongation rates as measured *in vitro* on FH1-FH2 fragments by (Scott et al., 2011) are shown on the right, as multiple of Fus1 elongation rate. **B.** DIC images ∼16h post starvation and GFP fluorescence images ∼8h post starvation of homothallic strains expressing C-terminally sfGFP-tagged chimeric formins with either (left) indicated FH2 domains and Fus1^FH1^ or (right) indicated FH2 domains and Cdc12^2FH1^. **C.** Percentage of cell pair fusion and lysis 24h after nitrogen removal in strains as in (B). Error bars are standard deviations. p-values (student’s t-test) of strains with Fus1^FH1^ chimeras are relative to Fus1^FH1^-Fus1^FH2^, p-values of strains with Cdc12^2FH1^ chimeras are relative to Cdc12^2FH1^-Fus1^FH2^. Bars are 5µm. **D.** ClustalO alignment of Cdc12 and Fus1 FH2 domains from *Schizosaccharomyces* species (Sp = *S. pombe*; Sc = *S. cryophilus*; So = *S. octosporus*; Sj = *S. japonicus*). The black boxes highlight mutated residues described in Figure 6.

**Figure S2 (related to Figure 6).**
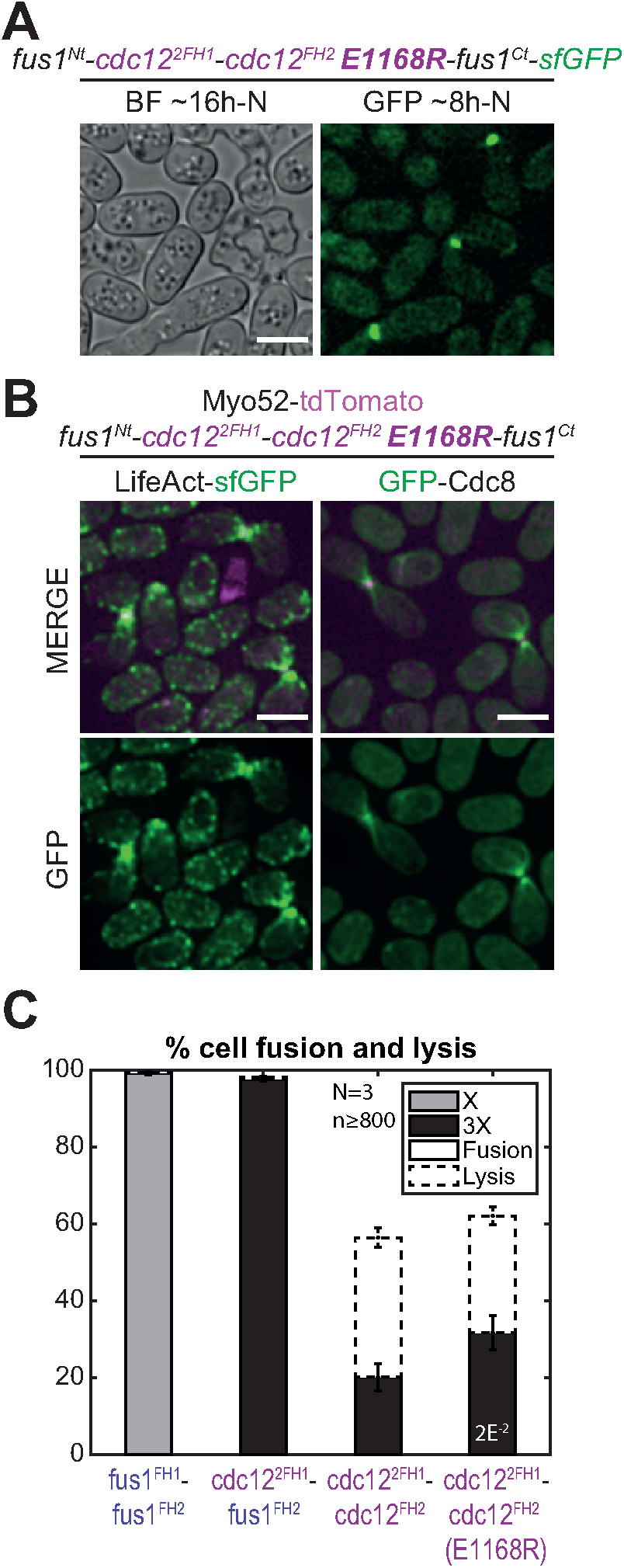
The complementary E1168R mutation in Cdc12 FH2 only slightly rescues the fusion defects observed with Cdc12 FH2. **A.** DIC images ∼16h post starvation and GFP fluorescence images ∼8h post starvation of homothallic strains expressing the formin chimera Cdc12^2FH1^-Cdc12E1168R^FH2^ tagged C-terminally with sfGFP. **B.** Merge and GFP fluorescence images ∼8h post starvation of Myo52-tdTomato and either (left) LifeAct-sfGFP or (right) GFP-Cdc8 in homothallic strains expressing the untagged formin chimera Cdc12^2FH1^-Cdc12E1168R^FH2^. **C.** Percentage of cell pair fusion and lysis 24h after nitrogen removal in strains as in (A) compared to the non-mutated controls. Error bars are standard deviations. p-value (student’s t-test) is relative to Cdc12^2FH1^-Cdc12^FH2^. Bars are 5µm.

**Figure S3.**
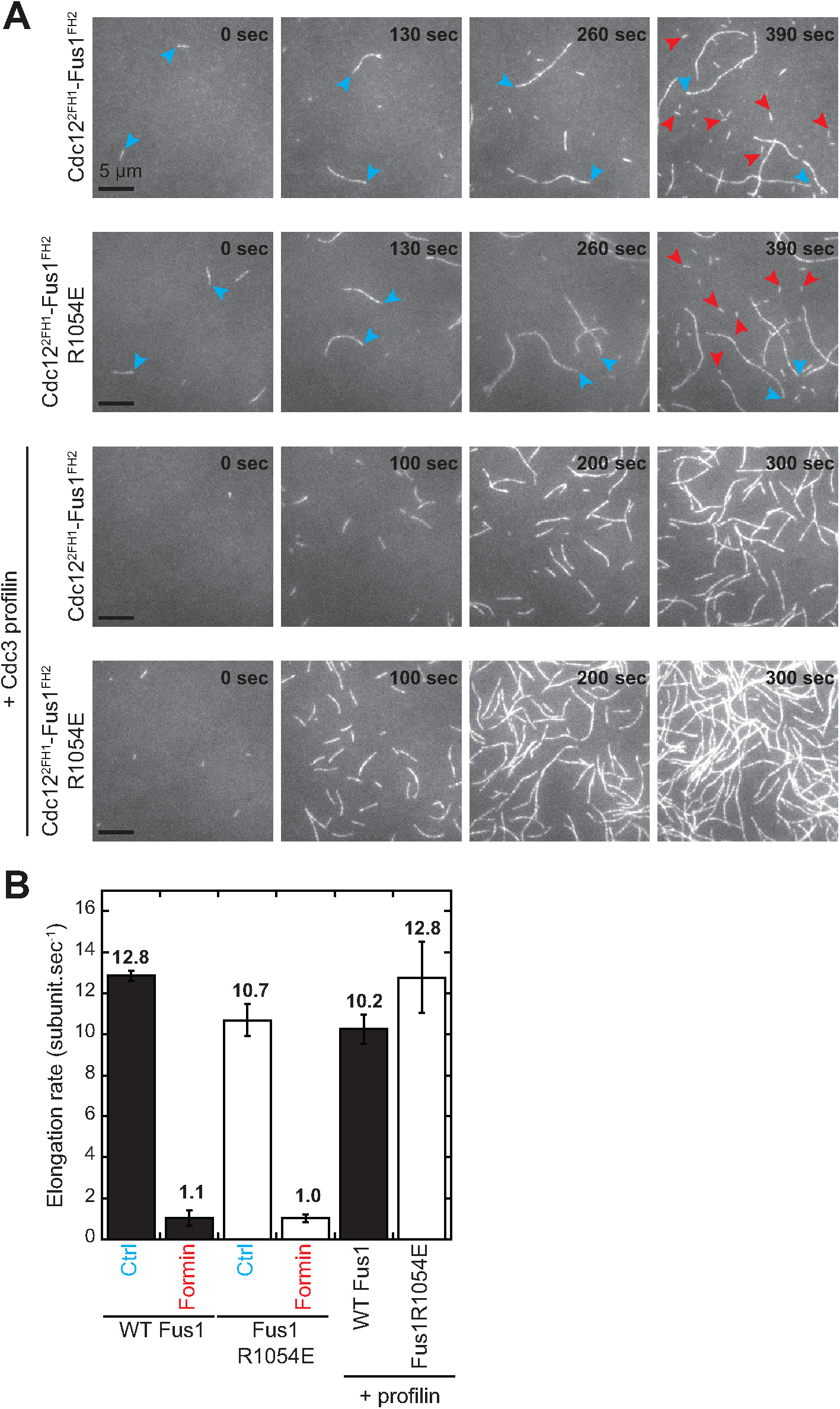
The actin filament elongation properties of Cdc12^2FH1^-Fus1^FH2^ are similar upon R1054E mutation. **A.** TIRF microscopy timelapse images of the polymerization of 1.5 µM Mg-ATP-actin (10% Alexa488-actin) with 500 pM of formin cdc12^2FH1^-Fus1^FH2^ (WT) or cdc12^2FH1^-Fus1_R1054E_^FH2^ (Mut) in the absence or presence of 2.5 µM profilin Cdc3. Blue arrowheads track two control (Ctrl) actin filaments elongating unbound to formin. Red arrows indicate formin-bound slow growing actin filaments. **B**. Average elongation rates of actin filaments in the conditions presented in panel (A). Error bars indicate standard errors.

**Movie S1 (related to Figure 1). Non fusing mating pairs expressing the Fus1N-Cdc12C chimera fail to coalesce their actin fusion foci into a single dot**

Time lapse images starting ∼4h post starvation of homothallic strains expressing the formin chimera Fus1N-Cdc12C tagged C-terminally with sfGFP (green) and Myo52-tdTomato (magenta). The movie shows three successful fusion events and one unsuccessful fusion (black arrowhead), in which the two partner cells form distant fusion foci (white arrowhead at 4h in the green channel and subsequent time points). Time is in hours:minutes. Bar is 5µm.

**Table S1:** Strain used in this study.

| GENOTYPE | FIGURES | STRAIN |
| --- | --- | --- |
| h90 myo52-tdTomato:natMX fus1-sfGFP:kanMX ura4- leu1-32 ade6-M216 | 1 2 3 4 6<br>S1 S2 | YSM3312 |
| h90 myo52-tdTomato:natMX fus1Δ::LEU2+ ura4-294:p<sup>fus1</sup>:fus1-sfGFP:ura4+ leu1-32 | 1 | YSM2498 |
| h90 myo52-tdTomato:natMX fus1Δ::LEU2+ ura4-294:p<sup>fus1</sup>:cdc12-sfGFP:ura4+ leu1-32 | 1 | YSM2502 |
| h90 myo52-tdTomato:natMX fus1Δ::LEU2+ ura4-294:p<sup>fus1</sup>:for3-sfGFP:ura4+ leu1-32 | 1 | YSM2500 |
| h90 myo52-tdTomato:natMX ura4+:p<sup>fus1</sup>:fus1N<sup>1-792</sup>-fus1C<sup>793-1372</sup>-sfGFP fus1Δ::hphMX ade6-M210 | 1 2 | YSM3942 |
| h90 myo52-tdTomato:natMX ura4+:p<sup>fus1</sup>:fus1N<sup>1-792</sup>-cdc12C<sup>888-1841</sup>-sfGFP fus1Δ::hphMX ade6-M210 | 1 2 | YSM3943 |
| h90 myo52-tdTomato:natMX ura4+:p<sup>fus1</sup>:fus1N<sup>1-792</sup>-for3C<sup>715-1461</sup>-sfGFP fus1Δ::hphMX ade6-M210 | 1 2 | YSM3944 |
| h90 myo52-tdTomato:natMX ura4+:p<sup>fus1</sup>:fus1N<sup>1-792</sup>-cdc12CΔolig<sup>888-1451</sup>-sfGFP fus1Δ::hphMX ade6-M210 | 1 | YSM3945 |
| h90 myo52-tdTomato:natMX fus1N<sup>1-792</sup>-fus1C<sup>793-1372</sup>-sfGFP:kanMX ura4-294 leu1-32 ade6-M210 | 2 | YSM3946 |
| h90 myo52-tdTomato:natMX fus1N<sup>1-792</sup>-cdc12C<sup>888-1841</sup>-sfGFP:kanMX ura4-294 leu1-32 ade6-M210 | 2 | YSM3947 |
| h90 myo52-tdTomato:natMX fus1N<sup>1-792</sup>-for3C<sup>715-1461</sup>-sfGFP:kanMX ura4-294 leu1-32 ade6-M210 | 2 | YSM3948 |
| h90 myo52-tdTomato:natMX fus1N<sup>1-792</sup>-fus1C<sup>793-1372</sup>-sfGFP:kanMX ura4-294 ade6-M210 | 2 | YSM3949 |
| h90 myo52-tdTomato:natMX fus1N<sup>1-792</sup>-cdc12C<sup>888-1841</sup>-sfGFP:kanMX ura4-294 ade6-M210 | 2 | YSM3950 |
| h90 myo52-tdTomato:natMX fus1N<sup>1-792</sup>-for3C<sup>715-1461</sup>-sfGFP:kanMX ura4-294 ade6-M210 | 2 | YSM3951 |
| h90 myo52-tdTomato:natMX ura4+:p<sup>fus1</sup>:fus1N<sup>1-792</sup>-fus1C<sup>793-1372</sup>-sfGFP fus1Δ::hphMX leu1-32 ade6-M210 | 2 | YSM3952 |
| h90 myo52-tdTomato:natMX ura4+:p<sup>fus1</sup>:fus1N<sup>1-792</sup>-cdc12C<sup>888-1841</sup>-sfGFP fus1Δ::hphMX leu1-32 ade6-M210 | 2 | YSM3953 |
| h90 myo52-tdTomato:natMX ura4+:p<sup>fus1</sup>:fus1N<sup>1-792</sup>-for3C<sup>715-1461</sup>-sfGFP fus1Δ::hphMX leu1-32 ade6-M210 | 2 | YSM3954 |
| h90 myo52-tdTomato:natMX fus1N<sup>1-792</sup>-fus1C<sup>793-1372</sup>-sfGFP:kanMX ura4+:p<sup>fus1</sup>:fus1N<sup>1-792</sup>-fus1C<sup>793-1372</sup>-sfGFP fus1Δ::hphMX leu1-32 ade6-M210 | 2 | YSM3955 |
| h90 myo52-tdTomato:natMX fus1N<sup>1-792</sup>-cdc12C<sup>888-1841</sup>-sfGFP:kanMX ura4+:p<sup>fus1</sup>:fus1N<sup>1-792</sup>-cdc12C<sup>888-1841</sup>-sfGFP fus1Δ::hphMX leu1-32 ade6-M210 | 2 | YSM3956 |
| h90 myo52-tdTomato:natMX fus1N<sup>1-792</sup>-for3C<sup>715-1461</sup>-sfGFP:kanMX ura4+:p<sup>fus1</sup>:fus1N<sup>1-792</sup>-for3C<sup>715-1461</sup>-sfGFP fus1Δ::hphMX leu1-32 ade6-M210 | 2 | YSM3957 |
| h90 myo52-tdTomato:natMX ura4+:p<sup>ste11</sup>:fus1-sfGFP fus1Δ::hphMX leu1-32 ade6-M210 | 2 | YSM3958 |
| h90 myo52-tdTomato:natMX ura4+:p<sup>pak2</sup>:fus1-sfGFP fus1Δ::hphMX leu1-32 ade6-M210 | 2 | YSM3959 |
| h90 myo52-tdTomato:natMX fus1<sup>Δ1278-1372</sup>-sfGFP:kanMX ura4-294 leu1-32 ade6-M210 | 3 | YSM3960 |
| h90 myo52-tdTomato:natMX fus1<sup>1-868</sup>-fus1<sup>792-1372</sup>-sfGFP:kanMX ura4-294 leu1-32 ade6-M210 | 4 | YSM3961 |
| h90 myo52-tdTomato:natMX fus1<sup>1-791</sup>-cdc12<sup>928-972</sup>-fus1<sup>869-1372</sup>-sfGFP:kanMX ura4-294 leu1-32 ade6-M210 | 4 | YSM3962 |
| h90 myo52-tdTomato:natMX fus1<sup>1-791</sup>-cdc12<sup>882-972</sup>-cdc12<sup>882-972</sup>-fus1<sup>869-1372</sup>-sfGFP:kanMX ura4-294 leu1-32 ade6-M210 | 4 6 S1 S2 | YSM3963 |
| h90 myo52-tdTomato:natMX ade6+:p<sup>act1</sup>:LifeAct-sfGFP:term<sup>ScADH1</sup>:bsdMX fus1:kanMX ura4-294 leu1-32 | 4 | YSM3964 |
| h90 myo52-tdTomato:natMX ade6+:p<sup>act1</sup>:LifeAct-sfGFP:term<sup>ScADH1</sup>:bsdMX fus1<sup>1-791</sup>-cdc12<sup>882-972</sup>-cdc12<sup>882-972</sup>-fus1<sup>869-1372</sup>:kanMX ura4-294 leu1-32 | 4 | YSM3965 |
| h90 myo52-tdTomato:natMX leu1-32:p<sup>nmt41</sup>:GFP-cdc8:ura4+ fus1:kanMX | 4 | YSM3966 |
| h90 myo52-tdTomato:natMX leu1-32:p<sup>nmt41</sup>:GFP-cdc8:ura4+ fus1<sup>1-791</sup>-cdc12<sup>882-972</sup>-cdc12<sup>882-972</sup>-fus1<sup>869-1372</sup>:kanMX | 4 | YSM3967 |
| h90 myo52-tdTomato:natMX fus1<sup>1-791</sup>-for3<sup>718-1265</sup>-fus1<sup>1278-1372</sup>-sfGFP:kanMX ura4-294 leu1-32 ade6-M210 | 5 | YSM3968 |
| h90 myo52-tdTomato:natMX fus1<sup>1-868</sup>-cdc12<sup>973-1390</sup>-fus1<sup>1278-1372</sup>-sfGFP:kanMX ura4-294 leu1-32 ade6-M210 | 5 6 S1 | YSM3969 |
| h90 myo52-tdTomato:natMX fus1<sup>1-791</sup>-cdc12<sup>928-1390</sup>-fus1<sup>1278-1372</sup>-sfGFP:kanMX ura4-294 leu1-32 ade6-M210 | 5 | YSM3970 |
| h90 myo52-tdTomato:natMX fus1<sup>1-791</sup>-cdc12<sup>882-1390</sup>-fus1<sup>1278-1372</sup>-sfGFP:kanMX ura4-294 leu1-32 ade6-M210 | 5 | YSM3971 |
| h90 myo52-tdTomato:natMX fus1<sup>1-792</sup>-cdc12<sup>882-972</sup>-cdc12<sup>882-1390</sup>-fus1<sup>1278-1372</sup>-sfGFP:kanMX ura4-294 leu1-32 ade6-M210 | 5 6 S1 S2 | YSM3972 |
| h90 myo52-tdTomato:natMX ade6+:p<sup>act1</sup>:LifeAct-sfGFP:term<sup>ScADH1</sup>:bsdMX fus1<sup>1-791-for3</sup><sup>718-1265-fus1<sup>1278-1372</sup></sup>:kanMX ura4-294 leu1-32 | 5 | YSM3973 |
| h90 myo52-tdTomato:natMX ade6+:p<sup>act1</sup>:LifeAct-sfGFP:term<sup>ScADH1</sup>:bsdMX fus1<sup>1-868-cdc12</sup><sup>973-1390-fus1<sup>1278-1372</sup></sup>:kanMX ura4-294 leu1-32 | 5 | YSM3974 |
| h90 myo52-tdTomato:natMX ade6+:p<sup>act1</sup>:LifeAct-sfGFP:term<sup>ScADH1</sup>:bsdMX fus1<sup>1-791-cdc12</sup><sup>928-1390-fus1<sup>1278-1372</sup></sup>:kanMX ura4-294 leu1-32 | 5 | YSM3975 |
| h90 myo52-tdTomato:natMX ade6+:p<sup>act1</sup>:LifeAct-sfGFP:term<sup>ScADH1</sup>:bsdMX fus1<sup>1-791-cdc12</sup><sup>882-972-cdc12<sup>882-1390-fus1<sup>1278-1372</sup></sup></sup>:kanMX ura4-294 leu1-32 | 5 | YSM3976 |
| h90 myo52-tdTomato:natMX leu1-32:p<sup>nmt41</sup>:GFP-cdc8:ura4+ fus1<sup>1-791-for3</sup><sup>718-1265-fus1<sup>1278-1372</sup></sup>:kanMX | 5 | YSM3977 |
| h90 myo52-tdTomato:natMX leu1-32:p<sup>nmt41</sup>:GFP-cdc8:ura4+ fus1<sup>1-868-cdc12</sup><sup>973-1390-fus1<sup>1278-1372</sup></sup>:kanMX | 5 | YSM3978 |
| h90 myo52-tdTomato:natMX leu1-32:p<sup>nmt41</sup>:GFP-cdc8:ura4+ fus1<sup>1-791-cdc12</sup><sup>928-1390-fus1<sup>1278-1372</sup></sup>:kanMX | 5 | YSM3979 |
| h90 myo52-tdTomato:natMX leu1-32:p<sup>nmt41</sup>:GFP-cdc8:ura4+ fus1<sup>1-791-cdc12</sup><sup>882-972-cdc12<sup>882-1390-fus1<sup>1278-1372</sup></sup></sup>:kanMX | 5 | YSM3980 |
| h90 myo52-tdTomato:natMX fus1<sup>1-868-Sj</sup><sup>fus1<sup>908-1317-fus1<sup>1278-1372</sup></sup></sup>-sfGFP:kanMX ura4-294 leu1-32 ade6-M210 | S1 | YSM3981 |
| h90 myo52-tdTomato:natMX fus1<sup>1-868-Sof</sup><sup>fus1<sup>857-1265-fus1<sup>1278-1372</sup></sup></sup>-sfGFP:kanMX ura4-294 leu1-32 ade6-M210 | S1 | YSM3982 |
| h90 myo52-tdTomato:natMX fus1<sup>1-792-cdc12</sup><sup>882-972-cdc12<sup>882-972-Sj</sup><sup>fus1<sup>908-1317-fus1<sup>1278-1372</sup></sup></sup></sup>-sfGFP:kanMX ura4-294 leu1-32 ade6-M210 | S1 | YSM3983 |
| h90 myo52-tdTomato:natMX fus1<sup>1-792-cdc12</sup><sup>882-972-cdc12<sup>882-972-Sof</sup><sup>fus1<sup>857-1265-fus1<sup>1278-1372</sup></sup></sup></sup>-sfGFP:kanMX ura4-294 leu1-32 ade6-M210 | S1 | YSM3984 |
| h90 myo52-tdTomato:natMX fus1(KEYTG935-939CARTD)-sfGFP:kanMX ura4-294 leu1-32 ade6-M216 | 6 | YSM3985 |
| h90 myo52-tdTomato:natMX fus1(L959K)-sfGFP:kanMX ura4-294 leu1-32 ade6-M216 | 6 | YSM3986 |
| h90 myo52-tdTomato:natMX fus1(EEVMEV1006-1011NGGDLVNS)-sfGFP:kanMX ura4-294 leu1-32 ade6-M216 | 6 | YSM3987 |
| h90 myo52-tdTomato:natMX fus1(R1054E)-sfGFP:kanMX ura4-294 leu1-32 ade6-M216 | 6 | YSM3988 |
| h90 myo52-tdTomato:natMX fus1(NHK1182-1184DPT)-sfGFP:kanMX ura4-294 leu1-32 ade6-M216 | 6 | YSM3989 |
| h90 myo52-tdTomato:natMX fus1<sup>1-792-cdc12</sup><sup>882-972-cdc12<sup>882-972-fus1<sup>869-1372</sup></sup></sup>(KEYTG935-939CARTD)-sfGFP:kanMX ura4-294 leu1-32 ade6-M210 | 6 | YSM3990 |
| h90 myo52-tdTomato:natMX fus1<sup>1-792-cdc12</sup><sup>882-972-cdc12<sup>882-972-fus1<sup>869-1372</sup></sup></sup>(L959K)-sfGFP:kanMX ura4-294 leu1-32 ade6-M210 | 6 | YSM3991 |
| h90 myo52-tdTomato:natMX fus1<sup>1-792-cdc12</sup><sup>882-972-cdc12<sup>882-972-fus1<sup>869-1372</sup></sup></sup>(EEVMEV1006-1011NGGDLVNS)-sfGFP:kanMX ura4-294 leu1-32 ade6-M210 | 6 | YSM3992 |
| h90 myo52-tdTomato:natMX fus1<sup>1-792-cdc12</sup><sup>882-972-cdc12<sup>882-972-fus1<sup>869-1372</sup></sup></sup>(R1054E)-sfGFP:kanMX ura4-294 leu1-32 ade6-M210 | 6 | YSM3993 |
| h90 myo52-tdTomato:natMX fus1<sup>1-792-cdc12</sup><sup>882-972-cdc12<sup>882-972-fus1<sup>869-1372</sup></sup></sup>(NHK1182-1184DPT)-sfGFP:kanMX ura4-294 leu1-32 ade6-M210 | 6 | YSM3994 |
| h90 myo52-tdTomato:natMX ade6+:p<sup>act1</sup>:LifeAct-sfGFP:term<sup>ScADH1</sup>:bsdMX fus1(R1054E):kanMX ura4-294 leu1-32 | 6 | YSM3995 |
| h90 myo52-tdTomato:natMX leu1-32:p<sup>nmt41</sup>:GFP-cdc8:ura4+ fus1(R1054E):kanMX | 6 | YSM3996 |
| h90 myo52-tdTomato:natMX ade6+:p<sup>act1</sup>:LifeAct-sfGFP:term<sup>ScADH1</sup>:bsdMX fus1<sup>1-792-cdc12</sup><sup>882-972-cdc12<sup>882-972-fus1<sup>869-1372</sup></sup></sup>(R1054E):kanMX ura4-294 leu1-32 | 6 | YSM3997 |
| h90 myo52-tdTomato:natMX leu1-32:p<sup>nmt41</sup>:GFP-cdc8:ura4+ fus1<sup>1-792-cdc12</sup><sup>882-972-cdc12<sup>882-972-fus1<sup>869-1372</sup></sup></sup>(R1054E):kanMX | 6 | YSM3998 |
| h90 myo52-tdTomato:natMX fus1<sup>1-792-cdc12</sup><sup>882-972-cdc12<sup>882-1390</sup></sup>(E1168R)-fus1<sup>1278-1372</sup>-sfGFP:kanMX ura4-294 leu1-32 ade6-M210 | S2 | YSM3999 |
| h90 myo52-tdTomato:natMX ade6+:p<sup>act1</sup>:LifeAct-sfGFP:term<sup>ScADH1</sup>:bsdMX fus1<sup>1-792-cdc12</sup><sup>882-972-cdc12<sup>882-1390</sup></sup>(E1168R)-fus1<sup>1278-1372</sup>:kanMX ura4-294 leu1-32 | S2 | YSM4000 |
| h90 myo52-tdTomato:natMX leu1-32:p<sup>nmt41</sup>:GFP-cdc8:ura4+ fus1<sup>1-792-cdc12</sup><sup>882-972-cdc12<sup>882-1390</sup></sup>(E1168R)-fus1<sup>1278-1372</sup>:kanMX | S2 | YSM4001 |

**Table S2:**
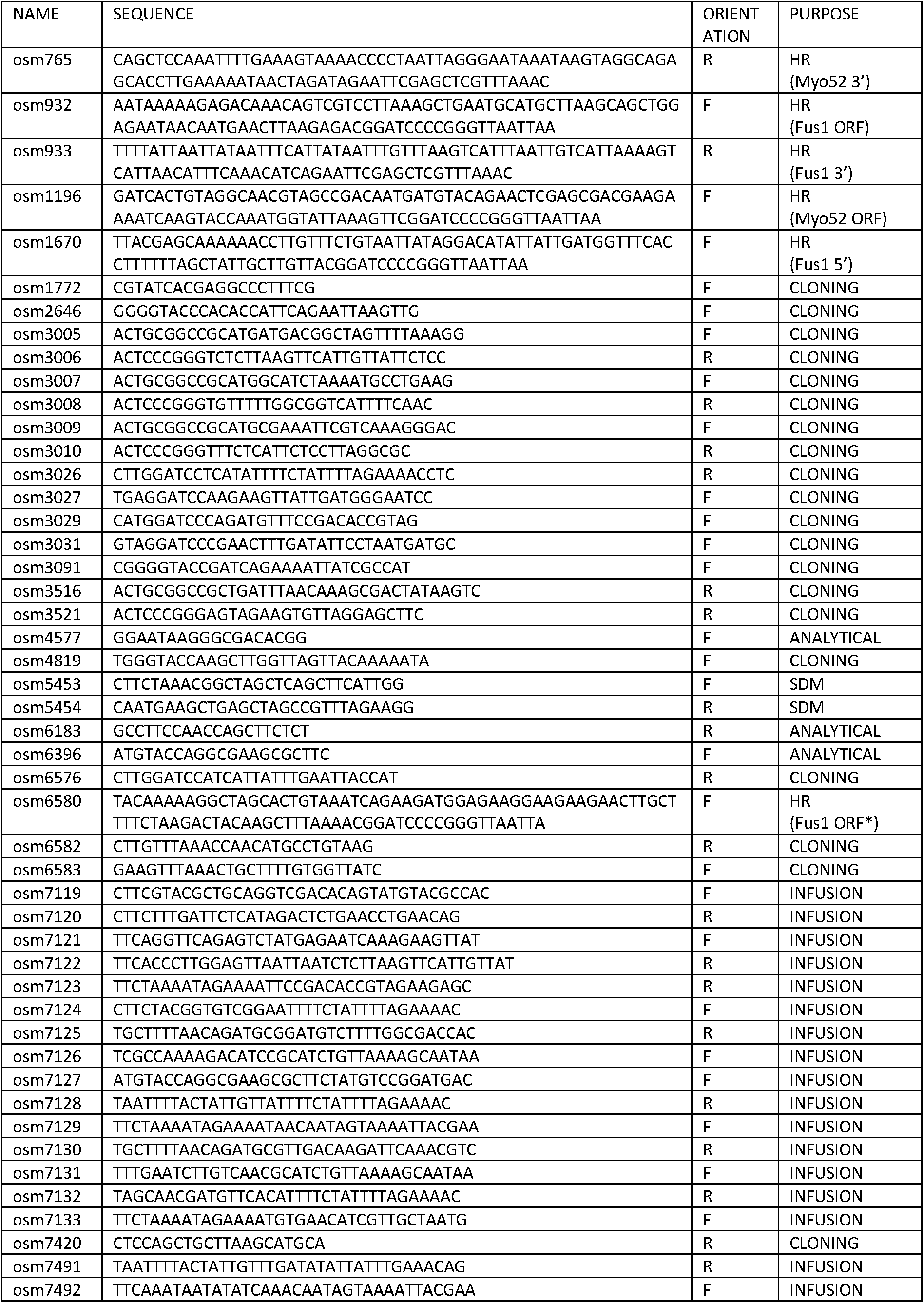
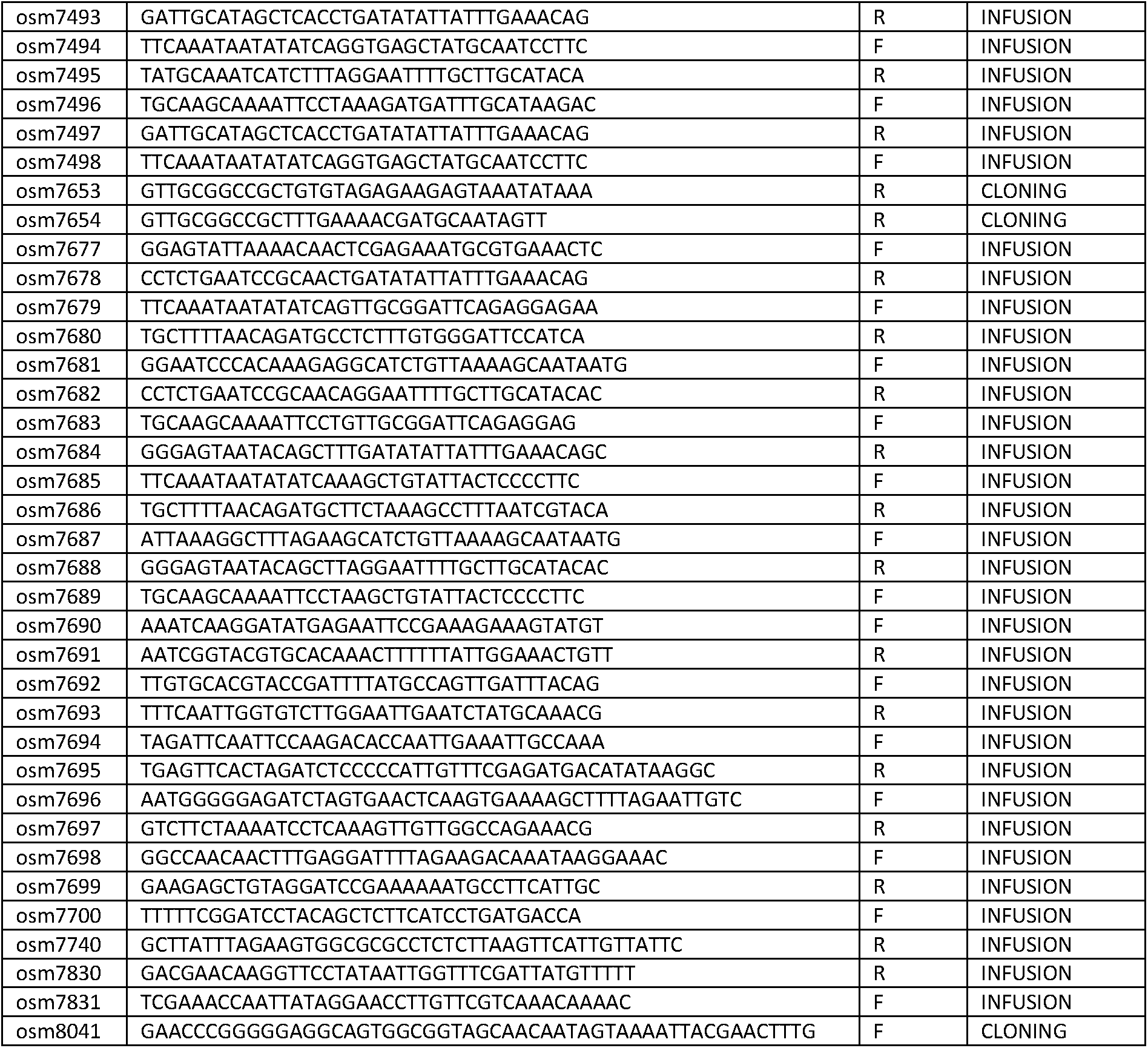
Primers used in this study. HR stands for homologous recombination in yeast (Bähler et al., 1998) and SDM for site directed mutagenesis

**Table S3:** Plasmids used in this study. For each plasmid, the column “obtained from” indicates how it was constructed, from restriction enzyme-based cloning or infusion, with the primers and restriction enzymes used. “WT” indicates that genomic DNA from a wildtype strain was used as template for PCR amplification.

| NAME | DESCRIPTION | OBTAINED FROM | USAGE |
| --- | --- | --- | --- |
| pAV133 | pUra4<sup>AfeI</sup> | (Vještica et al., 2020) | Single integration at ura4 |
| pAV530 | pUra4<sup>AfeI</sup>-p<sup>pak2</sup>-mCherry-SynZip3-term<sup>ScADH</sup> | Lab Stock, Derived from pAV133 | Single integration at ura4 |
| pSM685 | pFA6a-tdTomato-natMX | Lab Stock | template for PCR-based HR |
| pSM693 | pFA6a-hphMX | Lab Stock | template for PCR-based HR |
| pSM1538 | pFA6a-sfGFP-kanMX | Lab Stock | template for PCR-based HR |
| pSM1542 | pRIP82-p<sup>ste11</sup>-dVenus | Lab Stock | Multiple integration at ura4 |
| pSM1638 | pRIP-p<sup>fus1</sup>-sfGFP | Lab Stock | Multiple integration at ura4 |
| pSM1650 | pRIP-p<sup>fus1</sup>-fus1N-sfGFP | Regular cloning :<br>pSM1638<sup>NotI/BamHI</sup>+(WT<sup>osm3005-osm3026</sup>)<sup>NotI/BamHI</sup> | Multiple integration at ura4 |
| pSM1656 | pRIP-p<sup>fus1</sup>-fus1-sfGFP | Regular cloning :<br>pSM1638<sup>NotI/XmaI</sup>+(WT<sup>osm3005-osm3006</sup>)<sup>NotI/XmaI</sup> | Multiple integration at ura4 |
| pSM1657 | pRIP-p<sup>fus1</sup>-for3-sfGFP | Regular cloning :<br>pSM1638<sup>NotI/XmaI</sup>+(WT<sup>osm3007-osm3008</sup>)<sup>NotI/XmaI</sup> | Multiple integration at ura4 |
| pSM1658 | pRIP-p<sup>fus1</sup>-cdc12-sfGFP | Regular cloning :<br>pSM1638<sup>NotI/XmaI</sup>+(WT<sup>osm3009-osm3010</sup>)<sup>NotI/XmaI</sup> | Multiple integration at ura4 |
| pSM1659 | pRIP-p<sup>fus1</sup>-fus1N-fus1C-sfGFP | 3-point ligation cloning :<br>pSM1638<sup>NotI/XmaI</sup>+(WT<sup>osm3005-osm3026</sup>)<sup>NotI/BamHI</sup>+(WT<sup>osm3027-osm3006</sup>)<sup>BamHI/XmaI</sup> | Multiple integration at ura4 |
| pSM1660 | pRIP-p<sup>fus1</sup>-fus1N-for3C-sfGFP | 3-point ligation cloning :<br>pSM1638<sup>NotI/XmaI</sup>+(WT<sup>osm3005-osm3026</sup>)<sup>NotI/BamHI</sup>+(WT<sup>osm3029-osm3008</sup>)<sup>BamHI/XmaI</sup> | Multiple integration at ura4 |
| pSM1661 | pRIP-p<sup>fus1</sup>-fus1N-cdc12C-sfGFP | 3-point ligation cloning :<br>pSM1638<sup>NotI/XmaI</sup>+(WT<sup>osm3005-osm3026</sup>)<sup>NotI/BamHI</sup>+(WT<sup>osm3031-osm3010</sup>)<sup>BamHI/XmaI</sup> | Multiple integration at ura4 |
| pSM1823 | pRIP-p<sup>nmt41</sup>-sfGFP | Lab Stock | Multiple integration at ura4 |
| pSM1825 | pRIP-p<sup>fus1</sup>-fus1N-cdc12CΔolig-sfGFP | 3-point ligation cloning :<br>pSM1638<sup>NotI/XmaI</sup>+(WT<sup>osm3005-osm3026</sup>)<sup>NotI/BamHI</sup>+(WT<sup>osm3031-osm3521</sup>)<sup>BamHI/XmaI</sup> | Multiple integration at ura4 |
| pSM1826 | pRIP-p<sup>nmt41</sup>-fus1N-sfGFP | Subcloning :<br>pSM1650<sup>KpnI/NotI</sup>+pSM1823<sup>KpnI/NotI</sup> | Multiple integration at ura4 |
| pSM2229 | pUra4<sup>AfeI</sup>-p<sup>nmt41</sup>-fus1-sfGFP | Lab Stock, Derived from pAV133 | Single integration at ura4 |
| pSM2251 | pFA6a-fus1<sup>5'UTR</sup>-fus1(K879A)-sfGFP-kanMX-fus1<sup>3'UTR</sup> | Site-directed mutagenesis :<br>pSM2827<sup>osm5453/54354</sup> | Endogenous integration |
| pSM2478 | pUra4<sup>PmeI</sup>-p<sup>nmt41</sup>-fus1-sfGFP | 3-point ligation cloning :<br>pSM2229<sup>AatII/StuI</sup>+(pSM2229<sup>osm4577-osm6582</sup>)<sup>AatII/PmeI</sup>+(pSM2229<sup>osm6583-osm6183</sup>)<sup>PmeI/StuI</sup> | Single integration at ura4 |
| pSM2507 | pFA6a-fus1<sup>5'UTR</sup>-fus1<sup>Δ501-749</sup>-sfGFP-kanMX-fus1<sup>3'UTR</sup> | 3-point ligation cloning :<br>pSM2251<sup>Sall/PacI</sup>+(pSM2251<sup>osm1772-osm6576</sup>)<sup>Sall/BamHI</sup>+(pSM2229<sup>osm3031-osm3521</sup>)<sup>BamHI/PacI</sup> | Endogenous integration |
| pSM2594 | pUra4<sup>AfeI</sup>-p<sup>fus1</sup>-fus1N-fus1C-sfGFP | Subcloning :<br>pSM2478<sup>KpnI/SacI</sup>+pSM1659<sup>KpnI/SacI</sup> | Single integration at ura4 |
| pSM2595 | pUra4<sup>AfeI</sup>-p<sup>fus1</sup>-fus1N-cdc12C-sfGFP | Subcloning :<br>pSM2478<sup>KpnI/SacI</sup>+pSM1661<sup>KpnI/SacI</sup> | Single integration at ura4 |
| pSM2596 | pUra4<sup>AfeI</sup>-p<sup>fus1</sup>-fus1N-cdc12CΔolig-sfGFP | Subcloning :<br>pSM2478<sup>KpnI/SacI</sup>+pSM1825<sup>KpnI/SacI</sup> | Single integration at ura4 |
| pSM2600 | pUra4<sup>PmeI</sup>-p<sup>nmt1</sup>-fus1N-sfGFP | 3-point ligation cloning :<br>pSM2478<sup>KpnI/SacI</sup>+(pREP3x<sup>osm3091-osm3516</sup>)<sup>KpnI/NotI</sup><br>+pSM1826<sup>NotI/SacI</sup> | Single integration at<br>ura4 |
| pSM2602 | pUra4<sup>PmeI</sup>-p<sup>nmt1</sup>-fus1-sfGFP | Subcloning :<br>pSM2600<sup>NotI/XmaI</sup>+pSM1656<sup>NotI/XmaI</sup> | Single integration at<br>ura4 |
| pSM2604 | pUra4<sup>AfeI</sup>-p<sup>fus1</sup>-fus1N-for3C-sfGFP | Subcloning :<br>pSM2595<sup>Clal/MscI</sup>+pSM1660<sup>Clal/MscI</sup> | Single integration at<br>ura4 |
| pSM2621 | pFA6a-fus1<sup>5'UTR</sup>-fus1<sup>1-868-fus1<sup>792-1377</sup>-sfGFP-kanMX-fus1<sup>3'UTR</sup></sup> | Infusion cloning :<br>pSM2507<sup>Sall/PacI</sup>+WT<sup>osm7119-osm7120</sup>+WT<sup>osm7121-osm7122</sup> | Endogenous<br>integration |
| pSM2622 | pFA6a-fus1<sup>5'UTR</sup>-fus1<sup>1-791-cdc12<sup>882-1390</sup>-fus1<sup>1278-1372</sup>-sfGFP-kanMX-fus1<sup>3'UTR</sup></sup> | Infusion cloning :<br>pSM2507<sup>AfeI/PacI</sup>+WT<sup>osm7127-osm7128</sup>+ WT<sup>osm7129-osm7130</sup>+WT<sup>osm7131-osm7122</sup> | Endogenous<br>integration |
| pSM2631 | pFA6a-fus1<sup>5'UTR</sup>-fus1<sup>1-791-for3<sup>718-1265</sup>-fus1<sup>1278-1372</sup>-sfGFP-kanMX-fus1<sup>3'UTR</sup></sup> | Infusion cloning :<br>pSM2507<sup>Sall/PacI</sup>+WT<sup>osm6396-osm7123</sup>+ WT<sup>osm7124-osm7125</sup>+WT<sup>osm7126-osm7122</sup> | Endogenous<br>integration |
| pSM2632 | pFA6a-fus1<sup>5'UTR</sup>-fus1<sup>1-791-cdc12<sup>928-1390</sup>-fus1<sup>1278-1372</sup>-sfGFP-kanMX-fus1<sup>3'UTR</sup></sup> | Infusion cloning :<br>pSM2507<sup>AfeI/PacI</sup>+WT<sup>osm7127-osm7132</sup>+ WT<sup>osm7133-osm7130</sup>+WT<sup>osm7131-osm7122</sup> | Endogenous<br>integration |
| pSM2690 | pFA6a-fus1<sup>5'UTR</sup>-fus1N-fus1C-sfGFP-kanMX-fus1<sup>3'UTR</sup> | Subcloning :<br>pSM2251<sup>XhoI/XmaI</sup>+ pSM2594<sup>XhoI/XmaI</sup> | Endogenous<br>integration |
| pSM2691 | pFA6a-fus1<sup>5'UTR</sup>-fus1N-cdc12C-sfGFP-kanMX-fus1<sup>3'UTR</sup> | Subcloning :<br>pSM2251<sup>XhoI/XmaI</sup>+ pSM2595<sup>XhoI/XmaI</sup> | Endogenous<br>integration |
| pSM2692 | pFA6a-fus1<sup>5'UTR</sup>-fus1N-for3C-sfGFP-kanMX-fus1<sup>3'UTR</sup> | Subcloning :<br>pSM2251<sup>XhoI/XmaI</sup>+ pSM2604<sup>XhoI/XmaI</sup> | Endogenous<br>integration |
| pSM2699 | pFA6a-fus1<sup>5'UTR</sup>-fus1<sup>1-791-cdc12<sup>882-972</sup>-cdc12<sup>882-1390</sup>-fus1<sup>1278-1372</sup>-sfGFP-kanMX-fus1<sup>3'UTR</sup></sup> | Infusion cloning :<br>pSM2507<sup>AfeI/PacI</sup>+WT<sup>osm7127-osm7128</sup>+ WT<sup>osm7129-osm7491</sup>+WT<sup>osm7492-osm7130</sup>+WT<sup>osm7131-osm7122</sup> | Endogenous<br>integration |
| pSM2700 | pFA6a-fus1<sup>5'UTR</sup>-fus1<sup>1-791-cdc12<sup>882-972</sup>-cdc12<sup>882-972</sup>-fus1<sup>869-1372</sup>-sfGFP-kanMX-fus1<sup>3'UTR</sup></sup> | Infusion cloning :<br>pSM2507<sup>AfeI/PacI</sup>+WT<sup>osm7127-osm7128</sup>+ WT<sup>osm7129-osm7491</sup>+WT<sup>osm7492-osm7493</sup>+WT<sup>osm7494-osm7122</sup> | Endogenous<br>integration |
| pSM2701 | pFA6a-fus1<sup>5'UTR</sup>-fus1<sup>1-868-cdc12<sup>973-1390</sup>-fus1<sup>1278-1372</sup>-sfGFP-kanMX-fus1<sup>3'UTR</sup></sup> | Infusion cloning :<br>pSM2507<sup>AfeI/PacI</sup>+WT<sup>osm7127-osm7495</sup>+ WT<sup>osm7496-osm7130</sup>+WT<sup>osm7131-osm7122</sup> | Endogenous<br>integration |
| pSM2702 | pFA6a-fus1<sup>5'UTR</sup>-fus1<sup>1-791-cdc12<sup>928-972</sup>-fus1<sup>869-1372</sup>-sfGFP-kanMX-fus1<sup>3'UTR</sup></sup> | Infusion cloning :<br>pSM2507<sup>AfeI/PacI</sup>+WT<sup>osm7127-osm7132</sup>+ WT<sup>osm7133-osm7497</sup>+WT<sup>osm7498-osm7122</sup> | Endogenous<br>integration |
| pSM2827 | pFA6a-fus1<sup>5'UTR</sup>-fus1-sfGFP-kanMX-fus1<sup>3'UTR</sup> | Subcloning :<br>pSM2700<sup>EcoRI/NheI</sup>+ pSM2602<sup>EcoRI/NheI</sup> | Endogenous<br>integration |
| pSM2828 | pUra4<sup>PmeI</sup>-p<sup>ste11</sup>-fus1-sfGFP | Regular cloning :<br>pSM2602<sup>KpnI/NotI</sup>+(pSM1542<sup>osm2646-osm7653</sup>)<sup>KpnI/NotI</sup> | Single integration at<br>ura4 |
| pSM2829 | pUra4<sup>PmeI</sup>-p<sup>pak2</sup>-fus1-sfGFP | Regular cloning :<br>pSM2602<sup>KpnI/NotI</sup>+(pAV530<sup>osm4819-osm7654</sup>)<sup>KpnI/NotI</sup> | Single integration at<br>ura4 |
| pSM2844 | pFA6a-fus1<sup>5'UTR</sup>-fus1<sup>1-791-cdc12<sup>882-972</sup>-cdc12<sup>882-972</sup>-fus1<sup>869-1372</sup>(KEYTG935-939CARTD)-sfGFP-kanMX-fus1<sup>3'UTR</sup></sup> | Infusion cloning :<br>pSM2700<sup>EcoRI/PacI</sup>+pSM2700<sup>osm7690-osm7691</sup>+pSM2700<sup>osm7692-osm7122</sup> | Endogenous<br>integration |
| pSM2845 | pFA6a-fus1<sup>5'UTR</sup>-fus1<sup>1-791-cdc12<sup>882-972</sup>-cdc12<sup>882-972</sup>-fus1<sup>869-1372</sup>(L959K)-sfGFP-kanMX-fus1<sup>3'UTR</sup></sup> | Infusion cloning :<br>pSM2700<sup>EcoRI/PacI</sup>+pSM2700<sup>osm7690-osm7693</sup>+pSM2700<sup>osm7694-osm7122</sup> | Endogenous<br>integration |
| pSM2846 | pFA6a-fus1<sup>5'UTR</sup>-fus1<sup>1-791-cdc12<sup>882-972</sup>-cdc12<sup>882-972</sup>-fus1<sup>869-1372</sup>(EEVMEV1006-1011NGGDLVNS)-sfGFP-kanMX-fus1<sup>3'UTR</sup></sup> | Infusion cloning :<br>pSM2700<sup>EcoRI/PacI</sup>+pSM2700<sup>osm7690-osm7695</sup>+pSM2700<sup>osm7696-osm7122</sup> | Endogenous<br>integration |
| pSM2847 | pFA6a-fus1<sup>5'UTR</sup>-fus1<sup>1-791-cdc12<sup>882-972</sup>-cdc12<sup>882-972</sup>-fus1<sup>869</sup></sup> | Infusion cloning : | Endogenous<br>integration |
|  | 1372(R1054E)-sfGFP-kanMX-fus1<sup>3'</sup>UTR | pSM2700<sup>EcoRI/PacI</sup>+pSM2700<sup>osm7690-osm7697+pSM2700osm7698-osm7122</sup> |  |
| pSM2848 | pFA6a-fus1<sup>5'</sup>UTR-fus1<sup>1-791-cdc12<sup>882-972</sup>-cdc12<sup>882-972</sup>-fus1<sup>869-1372</sup>(NHK1182-1184DPT)-sfGFP-kanMX-fus1<sup>3'</sup>UTR</sup> | Infusion cloning :<br>pSM2700<sup>EcoRI/PacI</sup>+pSM2700<sup>osm7690-osm7699+pSM2700osm7700-osm7122</sup> | Endogenous integration |
| pSM2849 | pFA6a-fus1<sup>5'</sup>UTR-fus1(KEYTG935-939CARTD)-sfGFP-kanMX-fus1<sup>3'</sup>UTR | Infusion cloning :<br>pSM2700<sup>EcoRI/PacI</sup>+pSM2827<sup>osm7690-osm7691+pSM2700osm7692-osm7122</sup> | Endogenous integration |
| pSM2850 | pFA6a-fus1<sup>5'</sup>UTR-fus1(L959K)-sfGFP-kanMX-fus1<sup>3'</sup>UTR | Infusion cloning :<br>pSM2700<sup>EcoRI/PacI</sup>+pSM2827<sup>osm7690-osm7693+pSM2700osm7694-osm7122</sup> | Endogenous integration |
| pSM2851 | pFA6a-fus1<sup>5'</sup>UTR-fus1(EEVMEV1006-1011NGGDLVNS)-sfGFP-kanMX-fus1<sup>3'</sup>UTR | Infusion cloning :<br>pSM2700<sup>EcoRI/PacI</sup>+pSM2827<sup>osm7690-osm7695+pSM2700osm7696-osm7122</sup> | Endogenous integration |
| pSM2852 | pFA6a-fus1<sup>5'</sup>UTR-fus1(R1054E)-sfGFP-kanMX-fus1<sup>3'</sup>UTR | Infusion cloning :<br>pSM2700<sup>EcoRI/PacI</sup>+pSM2827<sup>osm7690-osm7697+pSM2700osm7698-osm7122</sup> | Endogenous integration |
| pSM2853 | pFA6a-fus1<sup>5'</sup>UTR-fus1(NHK1182-1184DPT)-sfGFP-kanMX-fus1<sup>3'</sup>UTR | Infusion cloning :<br>pSM2700<sup>EcoRI/PacI</sup>+pSM2827<sup>osm7690-osm7699+pSM2700osm7700-osm7122</sup> | Endogenous integration |
| pSM2854 | pFA6a-fus1<sup>5'</sup>UTR-fus1<sup>1-791-cdc12<sup>882-972</sup>-cdc12<sup>882-972</sup>-Sjfus1<sup>908-1317</sup>-fus1<sup>1278-1372</sup>-sfGFP-kanMX-fus1<sup>3'</sup>UTR</sup> | Infusion cloning :<br>pSM2700<sup>XhoI/PacI</sup>+pSM2700<sup>osm7677-osm7678+SjWT<sup>osm7679-osm7680</sup>+pSM2700osm7681-osm7122</sup> | Endogenous integration |
| pSM2855 | pFA6a-fus1<sup>5'</sup>UTR-fus1<sup>1-868-Sjfus1<sup>908-1317</sup>-fus1<sup>1278-1372</sup>-sfGFP-kanMX-fus1<sup>3'</sup>UTR</sup> | Infusion cloning :<br>pSM2700<sup>XhoI/PacI</sup>+pSM2701<sup>osm7677-osm7682+SjWT<sup>osm7683-osm7680</sup>+pSM2700osm7681-osm7122</sup> | Endogenous integration |
| pSM2856 | pFA6a-fus1<sup>5'</sup>UTR-fus1<sup>1-791-cdc12<sup>882-972</sup>-cdc12<sup>882-972</sup>-Sofus<sup>857-1265</sup>-fus1<sup>1278-1372</sup>-sfGFP-kanMX-fus1<sup>3'</sup>UTR</sup> | Infusion cloning :<br>pSM2700<sup>XhoI/PacI</sup>+pSM2700<sup>osm7677-osm7684+SjWT<sup>osm7685-osm7686</sup>+pSM2700osm7687-osm7122</sup> | Endogenous integration |
| pSM2857 | pFA6a-fus1<sup>5'</sup>UTR-fus1<sup>1-868-Sofus1<sup>857-1265</sup>-fus1<sup>1278-1372</sup>-sfGFP-kanMX-fus1<sup>3'</sup>UTR</sup> | Infusion cloning :<br>pSM2700<sup>XhoI/PacI</sup>+pSM2701<sup>osm7677-osm7688+SjWT<sup>osm7689-osm7686</sup>+pSM2700osm7687-osm7122</sup> | Endogenous integration |
| pSM2913 | pFA6a-fus1<sup>5'</sup>UTR-fus1-kanMX-fus1<sup>3'</sup>UTR | Infusion cloning :<br>pSM2827<sup>EcoRI/AscI</sup>+pSM2827<sup>osm7690-osm7740</sup> | Endogenous integration |
| pSM2914 | pFA6a-fus1<sup>5'</sup>UTR-fus1<sup>1-791-cdc12<sup>882-972</sup>-cdc12<sup>882-972</sup>-fus1<sup>869-1372</sup>-kanMX-fus1<sup>3'</sup>UTR</sup> | Infusion cloning :<br>pSM2827<sup>EcoRI/AscI</sup>+pSM2700<sup>osm7690-osm7740</sup> | Endogenous integration |
| pSM2915 | pFA6a-fus1<sup>5'</sup>UTR-fus1<sup>1-868-cdc12<sup>973-1390</sup>-fus1<sup>1278-1372</sup>-kanMX-fus1<sup>3'</sup>UTR</sup> | Infusion cloning :<br>pSM2827<sup>EcoRI/AscI</sup>+pSM2701<sup>osm7690-osm7740</sup> | Endogenous integration |
| pSM2916 | pFA6a-fus1<sup>5'</sup>UTR-fus1<sup>1-791-cdc12<sup>928-1390</sup>-fus1<sup>1278-1372</sup>-kanMX-fus1<sup>3'</sup>UTR</sup> | Infusion cloning :<br>pSM2827<sup>EcoRI/AscI</sup>+pSM2632<sup>osm7690-osm7740</sup> | Endogenous integration |
| pSM2917 | pFA6a-fus1<sup>5'</sup>UTR-fus1<sup>1-791-cdc12<sup>882-972</sup>-cdc12<sup>882-1390</sup>-fus1<sup>1278-1372</sup>-kanMX-fus1<sup>3'</sup>UTR</sup> | Infusion cloning :<br>pSM2827<sup>EcoRI/AscI</sup>+pSM2699<sup>osm7690-osm7740</sup> | Endogenous integration |
| pSM2918 | pFA6a-fus1<sup>5'</sup>UTR-fus1<sup>1-791-for3<sup>718-1265</sup>-fus1<sup>1278-1372</sup>-kanMX-fus1<sup>3'</sup>UTR</sup> | Infusion cloning :<br>pSM2827<sup>EcoRI/AscI</sup>+pSM2631<sup>osm7690-osm7740</sup> | Endogenous integration |
| pSM2924 | pFA6a-fus1<sup>5'</sup>UTR-fus1<sup>1-791-cdc12<sup>882-972</sup>-cdc12<sup>882-1390</sup>(E1168R)-fus1<sup>1278-1372</sup>-sfGFP-kanMX-fus1<sup>3'</sup>UTR</sup> | Infusion cloning :<br>pSM2700<sup>EcoRI/PacI</sup>+pSM2699<sup>osm7690-osm7830+pSM2699osm7831-osm7122</sup> | Endogenous integration |
| pSM2961 | pFA6a-fus1<sup>5'</sup>UTR-fus1(R1054E)-kanMX-fus1<sup>3'</sup>UTR | Infusion cloning :<br>pSM2827<sup>EcoRI/AscI</sup>+pSM2852<sup>osm7690-osm7740</sup> | Endogenous integration |
| pSM2962 | pFA6a-fus1<sup>5'</sup>UTR-fus1<sup>1-791-cdc12<sup>882-972</sup>-cdc12<sup>882-972</sup>-fus1<sup>869-1372</sup>(R1054E)-kanMX-fus1<sup>3'</sup>UTR</sup> | Infusion cloning :<br>pSM2827<sup>EcoRI/AscI</sup>+pSM2847<sup>osm7690-osm7740</sup> | Endogenous integration |
| pSM2963 | pFA6a-fus1<sup>5'</sup>UTR-fus1<sup>1-791-cdc12<sup>882-972</sup>-cdc12<sup>882-1390</sup>(E1168R)-fus1<sup>1278-1372</sup>-kanMX-fus1<sup>3'</sup>UTR</sup> | Infusion cloning :<br>pSM2827<sup>EcoRI/AscI</sup>+pSM2924<sup>osm7690-osm7740</sup> | Endogenous integration |
| KV1176 | pSNAP-fus1<sup>2FH1</sup>-fus1<sup>FH2</sup>-fus1<sup>Ct</sup> | Kovar Lab Stock | Bacterial expression |
| pSM2968 | pSNAP-cdc12<sup>2FH1</sup>-fus1<sup>FH2</sup>-fus1<sup>Ct</sup> | Regular cloning :<br>KV1176<sup>XmaI/NsiI</sup>+(pSM2914<sup>osm8041-osm7420</sup>)<sup>XmaI/NsiI</sup> | Bacterial expression |
| pSM2969 | pSNAP-cdc12<sup>2FH1</sup>-fus1<sup>FH2</sup>(R1054E)-<br>fus1<sup>Ct</sup> | Regular cloning :<br>KV1176<sup>XmaI/NsiI</sup>+(pSM2846<sup>osm8041-osm7420</sup>)<sup>XmaI/NsiI</sup> | Bacterial expression |

